# A high-density human mitochondrial proximity interaction network

**DOI:** 10.1101/2020.04.01.020479

**Authors:** Hana Antonicka, Zhen-Yuan Lin, Alexandre Janer, Woranontee Weraarpachai, Anne-Claude Gingras, Eric A. Shoubridge

## Abstract

We used BioID, a proximity-dependent biotinylation assay, to interrogate 100 mitochondrial baits from all mitochondrial sub-compartments to create a high resolution human mitochondrial proximity interaction network. We identified 1465 proteins, producing 15626 unique high confidence proximity interactions. Of these, 528 proteins were previously annotated as mitochondrial, nearly half of the mitochondrial proteome defined by Mitocarta 2.0. Bait-bait analysis showed a clear separation of mitochondrial compartments, and correlation analysis among preys across all baits allowed us to identify functional clusters involved in diverse mitochondrial functions, and to assign uncharacterized proteins to specific modules. We demonstrate that this analysis can assign isoforms of the same mitochondrial protein to different mitochondrial sub-compartments, and show that some proteins may have multiple cellular locations. Outer membrane baits showed specific proximity interactions with cytosolic proteins and proteins in other organellar membranes, suggesting specialization of proteins responsible for contact site formation between mitochondria and individual organelles. This proximity network will be a valuable resource for exploring the biology of uncharacterized mitochondrial proteins, the interactions of mitochondria with other cellular organelles, and will provide a framework to interpret alterations in sub-mitochondrial environments associated with mitochondrial disease.

**Bullet points:**

- We created a high resolution human mitochondrial protein proximity map using BioID
- Bait-bait analysis showed that the map has sub-compartment resolution and correlation analysis of preys identified functional clusters and assigned proteins to specific modules
- We identified isoforms of matrix and IMS proteins with multiple cellular localizations and an endonuclease that localizes to both the matrix and the OMM
- OMM baits showed specific interactions with non-mitochondrial proteins reflecting organellar contact sites and protein dual localization

## Introduction

Mitochondria house the machinery for a large number of essential cellular metabolic pathways including oxidative phosphorylation, heme and iron-sulfur cluster biosynthesis, the urea cycle, the tricarboxylic acid cycle, and beta oxidation. In addition, they have their own translation system for the polypeptides encoded in mitochondrial DNA, act as platforms for the innate immune system, buffer cellular calcium, and they control the release of proteins that initiate apoptosis. This myriad of functions is compartmentalized within the organelle to regulate mitochondrial metabolism in response to physiological signals and cellular demands. Although we have a good catalogue of mitochondrial proteins (Calvo et al., 2016) many remain poorly characterized or have completely unknown functions (Floyd et al., 2016; Pagliarini and Rutter, 2013). Mitochondrial diseases due to deficiencies in the oxidative phosphorylation machinery are amongst the most common inherited metabolic disorders (Frazier et al., 2019; Nunnari and Suomalainen, 2012). While nearly 300 causal genes have now been identified, the molecular basis for the extraordinary tissue specificity associated with these diseases remains an enduring mystery.

Mitochondria are double-membraned organelles, whose ultrastructure has been investigated for decades. The outer membrane (OMM), which serves as a platform for molecules involved in the regulation of innate immunity and for regulation of apoptosis, is permeable to metabolites and small proteins, and it forms close contacts with the endoplasmic reticulum (ER) for the exchange of lipids and calcium (Csordas et al., 2018). The impermeable inner membrane (IMM) is composed of invaginations called cristae that contain the complexes of the oxidative phosphorylation system. The intermembrane space (IMS) comprises both the space formed by the cristae invaginations, and the much smaller space between the inner boundary membrane and the OMM. A protein complex, the mitochondrial contact site and cristae organizing system (MICOS), bridges the neck of the cristae to contact sites in the OMM, and through that to the (ER) (Schorr and van der Laan, 2018). The matrix contains multitude of metabolic enzymes, as well as the components necessary for mitochondrial gene expression: the nucleoid, mitochondrial RNA granule, and the mitochondrial ribosome. Proteins are targeted to the different mitochondrial sub-compartments by specific import and sorting machineries (Wiedemann and Pfanner, 2017).

The first inventory of human mitochondrial proteins, Mitocarta, was created just over ten years ago by mass spectrometric analysis of mitochondria isolated from different human tissues (Pagliarini et al., 2008), and has since been updated to Mitocarta 2.0 (which includes 1158 proteins) using proteomic, machine-learning and data-mining approaches (Calvo et al., 2016). This database represents the most comprehensive, validated inventory of mitochondrial proteins. Using machine learning approaches, an Integrated Mitochondrial Protein Index (IMPI) created in Mitominer V4.0 (mitominer.mrc-mbu.cam.ac.uk) lists an additional 468 putative mitochondrial proteins, but most of these remain to be experimentally validated. The composition of individual mitochondrial sub-compartments has also been profiled by APEX, an engineered peroxidase used in a proximity-dependent biotinylation method to determine the composition of mitochondrial matrix (Rhee et al., 2013), the intermembrane space (Hung et al., 2014), and outer mitochondrial membrane proteomes (Hung et al., 2017). Although these datasets are core resources for mitochondrial biology, they do not provide information on functional relationships between individual proteins or the formation of protein complexes, nor do they allow one to assign uncharacterized proteins to a biological process.

Alternative approaches have provided added information on the organization of proteins within the mitochondria. Our group (Antonicka et al., 2017; Antonicka and Shoubridge, 2015; Janer et al., 2016) and others (Floyd et al., 2016) have defined, for mitochondrial proteins of interest, their interacting protein partners using affinity purification (or affinity enrichment)-mass spectrometry approaches. Large-scale investigations of protein-protein interactions have also been carried out by affinity purification followed by mass spectrometry using tagged, overexpressed baits (Huttlin et al., 2017; Malty et al., 2017). While these data are generally very useful in identifying soluble protein complexes, they are performed under conditions that may be sub-optimal for membrane proteins in organelles, and the analyses are carried out after cell lysis, which does not always represent the *in vivo* situation. In addition, they may not identify transient or weak interactions that do not survive the affinity purification protocols. More recently, chemical crosslinking coupled to mass spectrometry has shown promise in defining structural constraints amongst interacting mitochondrial proteins (Liu et al., 2018; Schweppe et al., 2017); however, the current coverage is sparse and does not provide a clear view of the overall mitochondrial organization.

We and others have previously employed, in a systematic manner, the proximity-dependent biotinylation assay, BioID, to reveal both the composition and the structural organization of membraneless organelles such as the centrosome (Gupta et al., 2015) and cytoplasmic RNA granules (P-bodies and stress granules (Youn et al., 2018)). BioID uses a mutated bacterial biotin ligase BirA*, fused to a protein of interest (bait), which biotinylates proteins with a solvent exposed lysine residue (preys) in the proximity of the bait (Roux et al., 2012). As with APEX, performing a single BioID experiment will not define protein complexes or structural organization. However, our previous work demonstrated that profiling an organelle with multiple baits yields spatial profiles of preys that can be used to reconstruct proximity relationships between preys (Gingras et al, 2019). This is because each bait will yield quantitative prey profiles that will preferentially label its closest (i.e. within ∼10nm) neighbors. While our published work focused on cytoplasmic membraneless organelles, we recently demonstrated that the approach could be applied to classical membrane-bound organelles as well to define a low-resolution structural organization of an entire human cells by profiling ∼200 subcellular localization markers by BioID (https://www.biorxiv.org/content/10.1101/796391v1). This study used 12 mitochondrial proteins as baits (including two mitochondrial signal sequences fused to BirA*), and while this was sufficient to define the mitochondrial space in the larger cell map, it did not provide deep insight into the sub-organellar organization.

Here we describe the creation of a high density human mitochondrial proximity network, created by interrogation of proximal interactors of 100 mitochondrial baits from all mitochondrial compartments. Using this technique, we show that we are able to quantitatively characterize the matrix and IMS environments, profile the proximity interactome in mitochondrial compartments and sub-compartments, assign preys to protein complexes, pathways or modules, demonstrate multiple localizations for isoforms of mitochondrial proteins, and to characterize the specific interactions of mitochondrial OMM proteins with other cellular organelles or protein complexes.

## Results

### Definition of the mitochondrial matrix and IMS BioID environments provides means to score enrichment of specific proximity interactions

BioID has been used for both defining the composition of structures including organelles, and to attempt to identify specific partners for baits of interest. The latter aspect is however challenged by the labeling of the environment simultaneously to that of specific proximal preys. While we expect that BioID will quantitively identify preys in the immediate neighborhood of a particular bait, prey identification is not necessarily directly proportional to their abundance as it also depends on both the number of solvent-exposed lysine residues and their individual reactivity. In very large datasets, specificity scoring across the entire dataset can be performed with for instance CompPASS (Sowa et al., 2009) or tools within ProHits-viz (Knight et al., 2017); however, this is not possible in datasets that only contain a few baits. A quantitative definition of subcellular and sub-organellar environments is therefore essential for the careful interpretation of BioID data. Previous definitions of “environments” have been performed by fusion of localization sequence tags, such as NLS (nuclear localization sequence) for the nucleus (Lambert et al., 2015), and a CAAX motif (Bagci et al., 2020) to study the specificity of Rho family GTPases at the plasma membrane. To define the matrix and IMS spaces, we used mitochondrial targeting sequences (MTS) to direct BirA* to these sub-compartments, similar to what was previously done using APEX (Hung et al., 2014; Rhee et al., 2013). To help mitigating potential biases due to the MTS sequences used, we sought to perform these experiments using different sequences.

BirA*-tagged MTS-baits were stably integrated into 293 Flp-In T-REx cells. We first validated localization to mitochondria by confocal microscopy using an anti-FLAG antibody (**Supplementary Fig. S1A**), followed by immunoblotting using the same antibody to assess the level of expression of the bait construct. In addition, we performed an immunoblot using an anti-biotin antibody to confirm that induction of the Bait-BirA* protein resulted in protein biotinylation. Each bait was analyzed by BioID in two to eight biological replicates (see Methods); reproducibility between the replicates calculated by Pearson correlation was between 0.89 and 0.98 (based on the spectral counts). High-confidence interactors for each bait were identified using SAINTexpress (Teo et al., 2014) by comparison to 48 controls (293 Flp-In T-REx cells alone to monitor endogenously biotinylated proteins or BirA*-GFP expressing cells to detect proteins that become promiscuously biotinylated), and we fixed the high-confidence SAINTexpress threshold at 1% Bayesian FDR.

To characterize the mitochondrial matrix environment, we fused BirA* to three of the most commonly used mitochondrial matrix targeting sequences; two from cytochrome *c* oxidase subunits COX4I1 (MTS-COX4) and COX8A (MTS-COX8), and one from ornithine carbamoyltransferase (OTC, MTS-OTC). While different MTS sequences share the same topology (amphipathic alpha helices), there is considerable variation in their length, but little or no conservation at the level of amino acid sequence. There is a vast literature on targeting heterologous proteins fused to an MTS to mitochondria, so we expected that all three constructs we used would identify largely the same group of preys, but we remained open to the possibility that different sequences might confer some specificity in targeting, an idea that to our knowledge has never been explored. We refer to the joint analysis of these MTS sequences as “Matrix-BirA*”. Matrix-BirA* overall recovered 267 high-confidence preys (**Fig. 1A, Supplementary Table S1 and S2**), with an overlap of 70% (187 preys) among the three baits. As expected, the comparison of Matrix-BirA* to either Mitocarta 2.0 or Gene Ontology Cellular Compartment (GOCC) revealed that Matrix-BirA* was enriched for proteins that were previously annotated as mitochondrial (249 preys/ 93%; **Supplementary Fig. 1B**), similar to the MTS-APEX (Rhee HW, Science 2013) where 94% of the detected 495 preys were annotated as mitochondrial, 233 of which were in common with Matrix BirA*. Sub-mitochondrial annotation was available for 142 Matrix-BirA* preys with 99% enrichment for mitochondrial matrix proteins; similarly, 97% of the MTS-APEX preys were specific to the matrix (**Supplementary Fig. S1B**). While the overall pairwise comparison between pairs of baits was high (average R^2^=0.76; range from 0.62 to 0.95), indicating that these three different MTS’s sample largely the same local environment and that the averaged Matrix-BirA* should be representative of the labeling propensity inside the matrix (see below), we noted that each of the baits induced preferential labeling (**Fig. 1B** and **Supplementary Fig. S1C**). The total number of spectra detected by each MTS-BirA* was nearly the same (14000-16000 total spectra), indicating that their expression levels were similar; however, comparison between MTS-OTC and MTS-COX8 (**Fig. 1B**) showed that MTS-OTC enriched proteins from the TCA cycle (GO:0006099) when compared to MTS-COX8. While this suggests that matrix MTS sequences might contain specific localization signals, the use of multiple targeting sequences allowed us to sample more proteins, and suggested that the average Matrix-BirA* is the most reliable reflection of the matrix environment.

**Figure 1.**
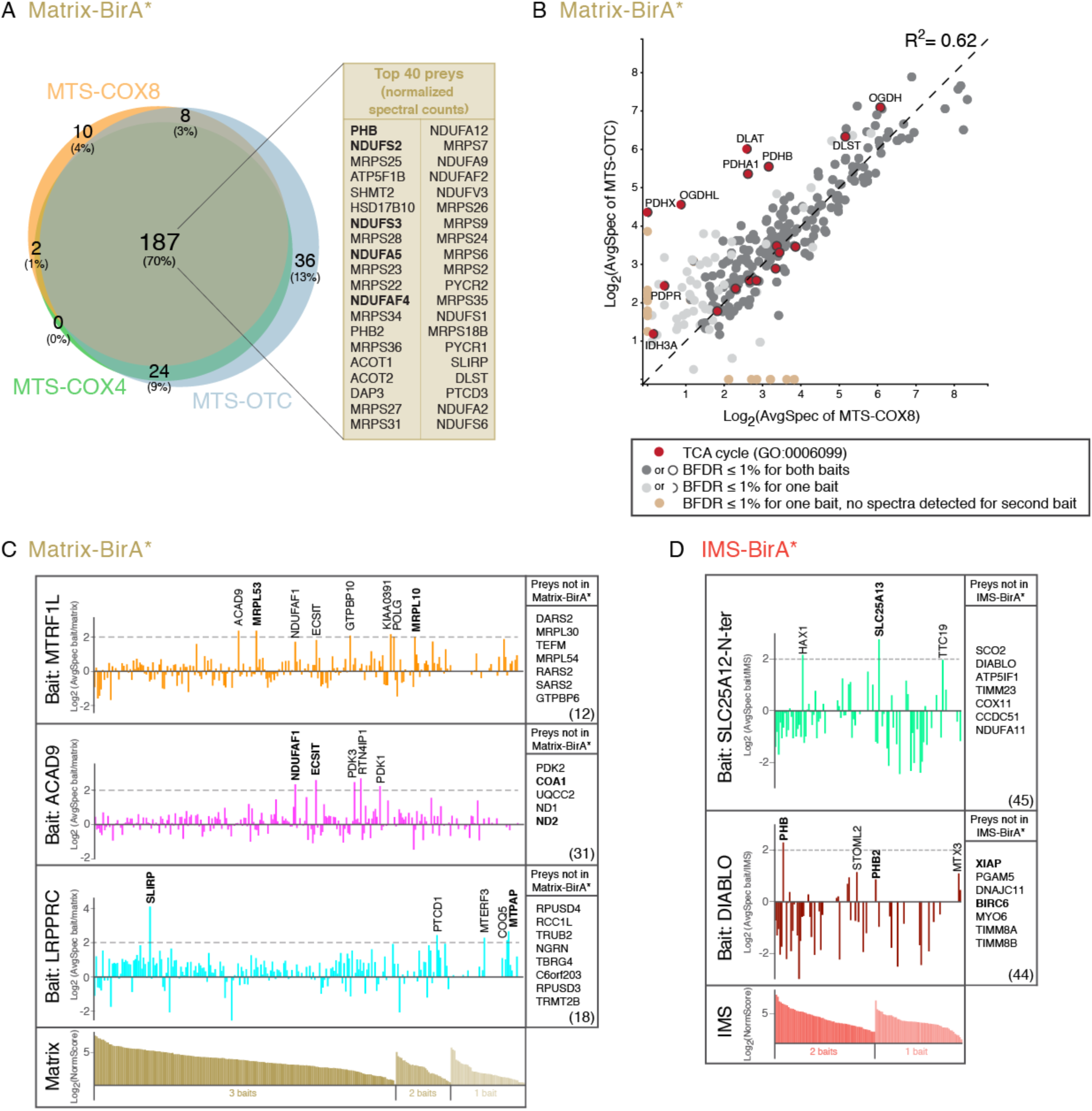
BioID definition of the mitochondrial matrix and IMS “environments”. **A.** Characterization of the mitochondrial matrix “environment”. Venn diagram of three Matrix-BirA* baits showing the most common set of interactors, the “environment”. Top 40 preys (based on normalized spectral counts) are indicated. The 5 preys in bold are common to both the matrix and IMS top 40 list; and are IMM proteins. **B.** Pairwise comparison of identified preys by MTS-OTC vs. MTS-COX8 to pinpoint the common and specific interactors for these baits (log_2_ transformed average spectral counts normalized to total abundance). The pairwise correlation coefficient, calculated from the untransformed spectral count matrix of the two conditions, is indicated. Proteins belonging to the TCA cycle (GO:0006099) are highlighted in red, with TCA cycle preys significantly enriched by MTS-OTC bait indicated by their gene name. TCA cycle proteins which passed BFDR for both baits are outlined with a full circle black border, proteins which passed BFDR only for one bait are indicated by a half circle border. Interactors in beige represent preys where no spectra were detected for the other bait, while interactors in light grey passed the BFDR≤1% only for one bait. Interactors common to both baits are in dark grey. **C.** Examples of the utilization of the matrix “environment” in identification of bait specific/enriched preys. The graphs represent log_2_ of the ratio between normalized spectral counts for each prey identified by the indicated bait and Matrix-BirA* score. Indicated preys for each bait represent the most enriched proximity interactors. Preys indicated in bold are previously described interactors. Each bait also interacted with a number of bait-specific preys that were not found as preys with BFDR≤1% by any of the three Matrix-BirA* baits (these are listed on the right in boxes; the number of proteins in this category is listed in parentheses; full list in **Supplementary Table S2**). The log_2_ of the normalized score for the 267 high-confidence preys (BFDR≤1%) identified by three mitochondrial Matrix-BirA* baits is shown at the bottom (brown bars) divided into three groups: preys identified by all three Matrix-BirA* baits (dark brown), two baits (medium brown) and by one bait only (light brown). The sorting order is based on the most abundant proteins seen across all three MTS baits, followed by those seen across two, followed by those seen in only one MTS. **D.** Examples of the utilization of the IMS “environment” in identification of bait specific/enriched preys, similar to (**C**). The graphs represent log_2_ of the ratio between normalized spectral counts for each prey identified by the indicated bait and IMS-BirA* score. Indicated preys for each bait represent the most enriched proximity interactors. Preys indicated in bold are previously described interactors. Each bait also interacted with a number of bait-specific preys which were not found as preys with BFDR≤1% by either of the two IMS-BirA* baits (these are listed on the right in boxes; the number of proteins in this category is listed in parentheses; full list in **Supplementary Table S2**). The log_2_ of the normalized score for the 120 high-confidence preys (BFDR≤1%) identified by two mitochondrial IMS-BirA* baits is shown at the bottom (red bars) divided into two groups: preys identified by two baits (red) and by one bait only (light red).

To test the usefulness of the Matrix-BirA* proteome in identifying specific proximity interactors for matrix baits, we analyzed it to score the prey enrichment for some well-characterized baits, namely LRPPRC, ACAD9 and MTRF1L. Briefly, BioID was performed in duplicates for each of these baits using the conditions described above, and scored with SAINTexpress, revealing lists of high-confidence proximity interactors ranging from 218 to 251 proteins (**Supplementary Table S2**), of which known members of protein complexes or direct interactors did not necessarily top the list. We next performed a fold-change enrichment of abundance of each of the interactors against the Matrix-BirA* to evaluate whether these known interactors were specifically enriched against the matrix “environment”. As shown in **Fig. 1C**, this was indeed the case: MTRF1L, a mitochondrial translation termination factor, specifically enriched MRPL53 and MRPL10, two mitochondrial ribosomal proteins present at the L7/L12 stalk, a site responsible for the recruitment of mitochondrial translation factors (Brown et al., 2014). Proximity interactors of ACAD9, a mitochondrial Complex I assembly factor, which is part of the MCIA complex responsible for ND2 module assembly (Formosa et al., 2018) were enriched for two other proteins of this complex (NDUFAF1 and ESCIT), as well as for ND2 protein itself and COA1, recently shown to be responsible for the translation of ND2 (Wang et al., 2020). Lastly, LRPPRC very strongly enriched SLIRP, with which it is known to form a stable complex (Sasarman et al., 2010) that is necessary for stabilization and polyadenylation of mitochondrial mRNAs (Chujo et al., 2012; Ruzzenente et al., 2012). Polyadenylation is performed by MTPAP, mitochondrial poly(A) polymerase, another proximity interactor of LRPPRC. Moreover, all three baits also showed a strong enrichment of additional proteins. For example: MTRF1L enriched the MCIA complex proteins (NDUFAF1, ACAD9, ECSIT); ACAD9 enriched three pyruvate dehydrogenase kinases (PDK1, PDK2, PDK3), whose activity results in inhibition of pyruvate dehydrogenase, and leads to decrease in oxidative phosphorylation; LRPPRC enriched proteins of mitochondrial RNA granule, more specifically with the pseudouridine-synthase module we previously reported (MTERF3, TRUB2, RPUSD3, RPUSD4, RCC1L, NGRN; (Antonicka et al., 2017)). All of these, previously undescribed associations suggest interesting research directions to explore.

The IMS environment was characterized using a “bipartite presequence” from two IMS proteins: OPA1 (MTS-OPA1) and apoptosis inducing factor AIFM1 (MTS-AIFM1). A “bipartite presequence” is a feature of a subset of IMS proteins that contain an MTS, followed by a cleavable stop-transfer sequence, which, upon cleavage, results in “soluble” IMS protein (Backes and Herrmann, 2017). Our IMS-BirA* baits identified 120 proximity interactors, of which 112 (93%) were annotated as mitochondrial proteins by either Mitocarta 2.0 or Gene Ontology Cellular Component (GOCC) (**Supplementary Fig. S1D**). These preys were highly enriched (97%) in IMS proteins, similar to IMS-APEX where 99% of preys were annotated as IMS proteins (Hung et al., 2014), but only 50 (42%) of the proteins identified in IMS-BirA* were in common with IMS-APEX. The pairwise comparison between our two IMS baits showed a high correlation of identified preys (R^2^=0.91); however, the overlap between the baits was only 53% (**Supplementary Fig. S1E-F**). This could result from biotinylation of preys prior to the cleavage of the pre-sequence, or possibly reflect targeting to a specific location in the IMM/IMS. MTS-AIFM1 for example preferentially labelled preys involved in protein targeting to mitochondria (GO:0006626, **Supplementary Fig. S1F**). One of these proteins, CHCHD4, has been previously reported to directly interact with full-length AIFM1 (Hangen et al., 2015; Reinhardt et al., 2020). The IMS compartment is organized on different principles than the matrix environment as it comprises the intra-cristae space, constrained by the narrow cristae necks formed by OPA1 and the MICOS complex, and the lumen between the OMM and the inner boundary membrane. As with the matrix proteome, we tested the usefulness of the IMS-BirA* proteome to identify specific proximity interactors by scoring for enrichment of proteins that had some prior characterization, namely SLC25A12 and DIABLO (**Fig. 1D, Supplementary Table S2**). SLC25A12, which codes for the mitochondrial aspartate/glutamate carrier and contains an approximately 300 amino acid long N-terminal domain in the IMS (Thangaratnarajah et al., 2014), enriched its paralogue SLC25A13, with which it has been shown to co-immunoprecipitate (Huttlin et al., 2017). Our analysis suggests that it may also associate with HAX1, an IMS protein which has been suggested to interact with the refoldase CLPB (Wortmann et al., 2015).

DIABLO, a “bipartite presequence” containing IMS protein, which is released in response to apoptotic signals, activates apoptosis by binding to inhibitors of apoptosis proteins XIAP and BIRC6 (Du et al., 2000), and both proteins were found as proximity interactors of DIABLO. Among the specifically enriched proximity interactors of DIABLO were three members of the SPFH family of scaffold proteins (PHB, STOML2 and PHB2). PHB was previously shown to immunoprecipitate DIABLO as well as XIAP to regulate apoptosis (Xu et al., 2016) and XL-MS investigations showed direct crosslinking between DIABLO and PHB2 (Koshiba and Kosako, 2019). Similar to the analysis of specificity enrichment for the matrix baits, the analysis of specificity of IMS baits (or IMM baits facing IMS) revealed the usefulness of the IMS-BirA* score as a quantitative assessment of the IMS “environment”.

These analyses demonstrate the utility of the Matrix-BirA* and IMS-BirA* resource to help interpret BioID experiments, and the illustrative examples show how the identification of specific proximity interactions for individual baits can be useful in compiling and narrowing down a list of candidate proteins for mechanistic investigation. Both Matrix-BirA* and IMS-BirA* scores are readily available in **Supplementary Table S2**.

### A comprehensive exploration of mitochondria by BioID

While the results above demonstrate that establishing the environment of mitochondrial sub-compartments facilitates interpretation of BioID data, we next asked whether profiling BioID by multiple baits for each of the mitochondrial sub-compartments would shed additional light on the spatial organization of proteins within mitochondrial domains. To do so, we selected ∼10% of human mitochondrial proteome (based on Mitocarta 2.0) as baits (**Fig. 2A**). We used two main criteria for bait selection. First, we wanted to interrogate all sub-mitochondrial compartments, including OMM, IMS, IMM facing both the IMS and the matrix, and the matrix space (**Fig. 2B**). Second, we selected baits that we thought were most likely to illuminate new aspects of mitochondrial biology that may not have been discovered or were poorly understood, with the hope of being able to assign preys of unknown function to biochemical pathways or protein complexes. Thus, we did not for instance choose as baits structural subunits of the oxidative phosphorylation complexes, the crystal structures of which have already been solved in several prokaryotic and eukaryotic systems (e.g. (Zhu et al., 2016; Zong et al., 2018)).

**Figure 2.**
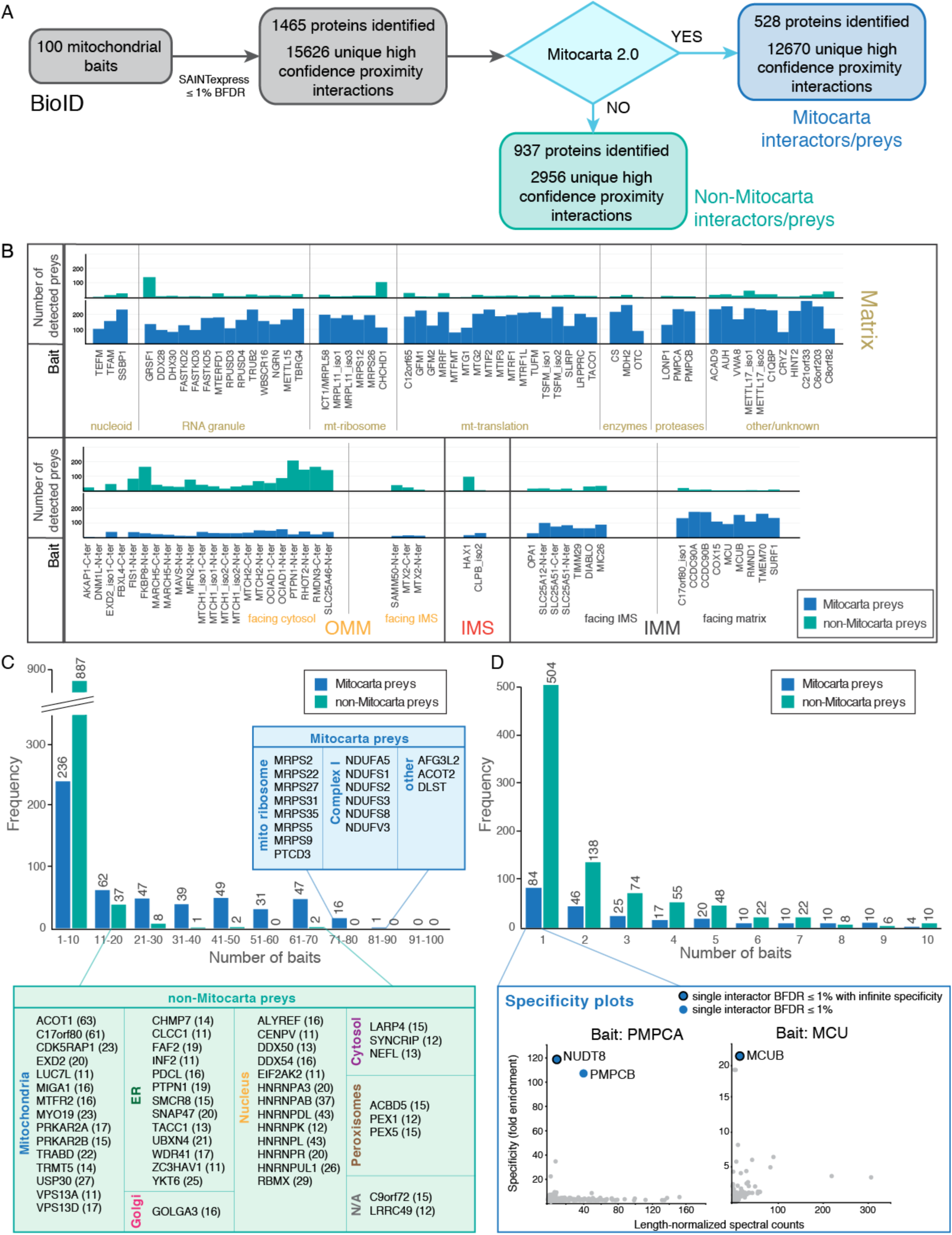
A comprehensive exploration of mitochondria by BioID. **A.** Project summary for the creation of mitochondrial interaction network using the proximity biotinylation assay (BioID) with 100 mitochondrial baits. **B.** A list of mitochondrial baits used in BioID from four individual mitochondrial sub-compartments and the number of identified preys for each bait, based on their previous annotation by Mitocarta 2.0 (Calvo et al., 2016). Preys were classified as Mitocarta or non-Mitocarta preys if annotated as mitochondrial, or not, respectively. Membrane spanning baits are divided into two groups depending on the orientation of the BirA*-tag. **C, D**. Frequency distribution of individual Mitocarta or non-Mitocarta preys across all the baits analysed. In (**D**) close-up of frequency of prey detection by 1-10 baits. (**C**) The lists of the most commonly identified Mitocarta preys (by more than 70 baits) and non-Mitocarta preys (by more than 10 baits). HPA protein localization and proximity map of the human cell were used to determine the cellular localization of non-Mitocarta preys. The number in parenthesis following the prey name represents the number of baits interacting with the identified prey. (**D**) Specificity plot examples of two baits: matrix bait PMPCA and IMM bait MCU indicating the specific interaction with preys identified as single interactors in our dataset. Infinite specificity signifies that no spectrum was detected for that prey in any other sample analyzed.

While we aimed at profiling a number of baits roughly proportional to the proteome of each compartment, the set of baits we choose was slightly weighted to those predicted to be targeted to the matrix, as this compartment likely contains the most substructures, including the nucleoid, the mitochondrial ribosome and other components of the translation apparatus, and the mitochondrial RNA granule (**Fig. 2B**). It was difficult to find soluble IMS baits that would properly localize to IMS with a BirA* tag. Many IMS proteins belong to the twin-CX_9_C motif-containing family, and the presence of CX_9_C motif at the C-terminus of the protein is essential for their import via Mia40 (CHCHD4) pathway (Chacinska et al., 2004), precluding their C-terminal tagging (the N-terminus also cannot be used because the mitochondrial targeting sequence, MTS, is located there). Nevertheless, we were able to explore the IMS space using two soluble IMS proteins (CLPB and HAX1) and several proteins of the inner mitochondrial membrane (IMM) with the BirA*-tag facing the IMS. For OMM proteins, the position of the tag was either inferred from literature, based on the protein structure profiling using a transmembrane prediction program TMHMM (http://www.cbs.dtu.dk/services/TMHMM/), or tagging of both ends was tested. Finally, nine OMM proteins were tagged at the N-terminus, four at the C-terminus, and 6 proteins were tagged at both N- and C-terminus. Those baits, in which both the N-and C-termini faced the cytoplasm, might in principle identify a different prey set, which could give us clues as to function. There is very little data on isoforms in the mitochondrial proteome, and we profiled two different isoforms for four genes to begin exploring this aspect of mitochondrial biology (**Supplementary Table S3, S8**).

As described above for analysis of the matrix and IMS environments, BirA*-tagged baits were stably integrated into 293 Flp-In T-REx cells and analyzed by BioID in biological duplicates (see Methods). The same quality controls were performed, and the corresponding confocal microscopy images are shown on http://prohits-web.lunenfeld.ca (currently password-protected; see Methods). A pairwise comparison between the two biological replicates for the same bait averaged R^2^=0.92 (range 0.60-0.99). High-confidence interactors for each bait were identified using SAINTexpress (Teo et al., 2014), as mentioned above. After fixing the high-confidence SAINTexpress threshold at 1% Bayesian FDR, 1465 proteins and 15626 unique proximity interactions were detected (**Fig. 2A, Supplementary Table S4**). Of these, 528 proteins (producing 12,670 unique high-confidence proximity interactions) were annotated as mitochondrial by Mitocarta 2.0 (here and throughout the paper referred to as “Mitocarta interactors/preys”), which corresponded to 46% of the mitochondrial proteome defined by Mitocarta 2.0. The total number of detected preys per bait only moderately correlated with the expression level of the individual baits (r=0.4915, P<0.0001) (**Supplementary Fig. S2A**).

As expected, matrix baits detected predominantly mitochondrial preys (**Fig. 2B, Supplementary Fig. S2A**), while outer mitochondrial membrane (OMM) baits detected a large number of proteins not annotated in Mitocarta (“non-Mitocarta preys”) (for details of OMM interactors see below). Only three baits that were not targeted to the OMM, namely HAX1 in the IMS, and the matrix proteins GRSF1 and CHCHD1, identified a significant number of non-Mitocarta preys, suggesting that they may be targeted to more than one compartment. GRSF1 has two isoforms: mitochondrial matrix GRSF1, which binds G-rich RNAs transcribed from the light strand of mtDNA (Antonicka et al., 2013), and another shorter isoform, present in some cells, that can be detected on immunoblots, and disappears in protease protection assays on isolated mitochondria ((Jourdain et al., 2013) and unpublished data). The latter could be translated from a downstream in frame ATG, and our data suggest that this isoform localizes to the nucleolus, where the BirA*-tagged bait biotinylates large cytosolic ribosomal subunit components (**Supplementary Fig. S2B**). CHCHD1 is a subunit of mitochondrial ribosome, and its role outside mitochondria is unknown. We show that it also recovers a number of nucleolar proteins, but also proteins of the nuclear envelope (**Supplementary Fig. S2B**). HAX1 has been shown to function both within mitochondria as well as outside, and has been suggested to express at least 6 isoforms (Lees et al., 2008), and our data are consistent with multiple predicted localizations for this protein.

In total, BioID analysis identified as proximity interactors 528 proteins previously annotated as mitochondrial proteins and 937 proteins not annotated in Mitocarta 2.0 (**Fig. 2A**). Of these, 17 Mitocarta preys that were identified as proximity interactors for more than 70 different baits (**Fig. 2C**), were proteins in either the matrix or IMM facing the matrix, and all are also part of the matrix “environment”. The majority of non-Mitocarta interactors (95%) were detected by fewer than ten baits (**Fig. 2C**), with 54% of non-Mitocarta preys detected only once (**Fig. 2D**). The remaining 50 non-Mitocarta preys detected by more than ten baits, could either be novel mitochondrial proteins, proteins present in a different cellular compartment which is in contact with mitochondria, or potentially artefactual proteins. All 50 of these preys were compared to Human Protein Atlas cellular localization data, in which proteins have been profiled by immunofluorescence using antibodies against the endogenous proteins and with our proximity map of a human cell (https://www.biorxiv.org/content/10.1101/796391v1). At least 15 proteins (ACOT1, C17orf80, CDK5RAP1, EXD2, LUC7L, MIGA1, MTFR2, MYO19, PRKAR2A, PRKAR2B, TRABD, TRMT5, USP30, VPS13A, VPS13D) were shown or suggested to localize to mitochondria by at least one of the profiles (**Fig. 2C**). Additionally, studies focusing on CDK5RAP1, MIGA1, MTFR2, MYO19, USP30 or VPS13A showed that these proteins either localize or tether to mitochondria (Bingol et al., 2014; Kumar et al., 2018; Lu et al., 2019; Shneyer et al., 2016; Yamamoto et al., 2019; Zhang et al., 2016). EXD2, an endonuclease, was originally described as a nuclear protein with a role in double-stranded DNA breaks (Broderick et al., 2016), but was later shown to localize to mitochondria (Silva et al., 2018) (Hensen et al., 2018); however, with a discrepancy in its sub-mitochondrial localization, a puzzle that we were able to resolve using BioID (see below). This analysis suggests that at least some of these proteins may be missing from annotation in Mitocarta 2.0, and shows the value of our dataset in the detection of novel mitochondrial proteins.

Interestingly, 84 Mitocarta preys (16%) were scored as high-confidence proximity interactors for a single bait (**Fig. 2D, Supplementary Table S5)**, and 14 of these 84 preys were never seen by any of the other baits in our dataset (infinite specificity). While most of these were in relatively low abundance, even with the bait with which they scored (median number of spectra detected was six), **Fig. 2D** and **Supplemental Fig. S2C** show some examples of these unique proximity interactions using “prey specificity plots”, which represent the specificity of proximity interaction between the identified prey and the bait of interest compared to all the other baits in the dataset (using spectral count abundance measurements). We notably show that PMPCA, as a bait, strongly and specifically enriched PMPCB (**Fig. 2D**). These two proteins form a complex, the mitochondrial processing peptidase, and both are required for its activity (Saavedra-Alanis et al., 1994). Another example is the proximity interaction between MCU (mitochondrial calcium uniporter) and MCUB, which form the complex responsible for Ca^2+^ import into mitochondrial matrix (Raffaello et al., 2013). Specificity plots for ACAD9 and SLC25A12 (**Supplemental Fig. S2C**) also show single proximity interactors, in addition to the same specific interactors shown in **Fig. 1C and D**, highlighting the fact that the Matrix-BirA* and IMS-BirA* datasets are good proxies for the entire dataset.

Overall our global analysis provides a rich, high quality dataset of proximity interactions with baits representing all mitochondrial compartments and sub-compartments that can be mined to explore functional relationships, both within the organelle itself and between the organelle and other cellular compartments.

### Bait self-organization reveals distinct matrix clusters

To globally analyze the data, we next compared the similarity in profiles of all the baits in our dataset using a Jaccard similarity coefficient, followed by hierarchical clustering. As expected, baits predicted to profile the same space have a higher Jaccard index with one another and a lower Jaccard distance score. **Fig. 3** shows the bait-bait heatmap of Jaccard distance score of the baits in our dataset. OMM baits, which are separated from the matrix baits by one, or even two membranes (if the tag is facing the cytosol) clearly profiled different preys than the matrix baits, as the Jaccard distance score was between 0.9–1.0. IMM baits, although in general clustering with the matrix baits, could be easily separated based on the orientation of the BirA* tag: if the tag was facing the matrix, the Jaccard distance to the matrix baits was <0.7 and to the OMM baits was >0.8. Most baits separately tagged at the N-and C-terminus appeared in the same cluster, with the exception of MTCH1 and MTCH2, which, although they clustered near each other, were clearly distinct. Likewise, the isoforms we used as baits clustered together, with the exception of TSFM-iso1and -iso2, which appeared in two distinct clusters, but in the same matrix compartment.

**Figure 3.**
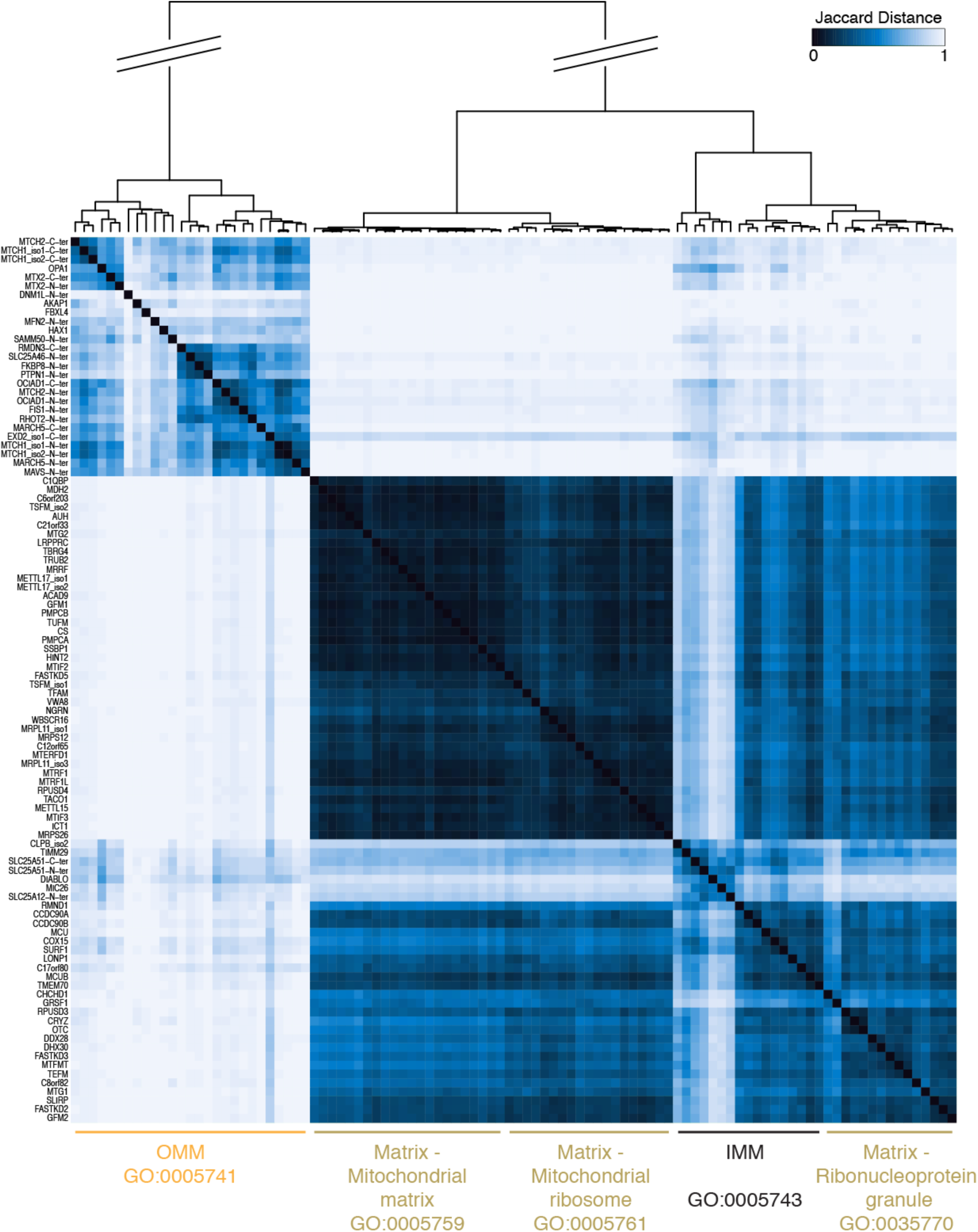
Bait self-organization reveals distinct matrix clusters. A bait-bait correlation heatmap created using the Jaccard distance score shows a clear separation between individual mitochondrial sub-compartments, as well as between distinct clusters within the mitochondrial matrix, highlighted by the dendrogram on the top of the heatmap. The Jaccard distance is defined as 1 – J(A,B), where J(A,B) is the Jaccard index, which defines the similarity between sample sets; the closest baits show the lowest distance score. Bait names are indicated on the left of the heatmap. Baits belonging to individual clusters were profiled in Panther, and the identified GO terms Cellular Compartment are indicated below the heatmap.

Interestingly, even by this relatively crude similarity measure, matrix baits found in different sub-compartments formed distinct clusters, which were annotated by their GOCC terms (**Fig. 3**). The baits in the mitochondrial matrix sub-cluster showed a high correlation (Jaccard distance score <0.17), and included mostly enzymes, proteases and proteins associated with mitochondrial translation and assembly of the oxidative phosphorylation complexes with different functions. The ribonucleoprotein granule cluster included GRSF1, FASTKD2, DDX28 and DHX30, proteins that we have previously shown to co-immunoprecipitate (Antonicka et al., 2013) and that are molecular markers of the mitochondrial RNA granule. The mitochondrial RNA granule is also a hub for mitoribosome biogenesis, and several proteins implicated in this process clustered in the mitochondrial ribosome cluster (MTERFD1, NGRN, WBSCR16 (RCC1L), RPUSD4, METTL15). This indicates that the approach has compartment and sub-compartment spatial resolution. To further investigate this, we examined pairs of baits within a cluster, across different clusters of the matrix, and across different sub-compartments by performing Pearson correlation analysis of the spectral counts of the preys identified (**Supplementary Fig. S3A**). The average Pearson correlation across all matrix baits was 0.71, across the OMM baits was 0.57, and between OMM and matrix was 0.28, distinctly separating baits at the extremities of mitochondria. Examples of the representative pairwise comparisons between matrix-matrix, OMM-OMM and matrix-OMM baits are shown in **Supplementary Fig. S3B-D**.

As expected, though the correlation coefficients are modest, OMM baits correlated more strongly with IMM baits facing the IMS (r=0.42), compared to r=0.30 between OMM and IMM facing matrix, while matrix baits correlated more with IMM facing the matrix (r=0.54) compared to r=0.31 between matrix and IMM facing IMS (**Supplementary Fig. S3A**). The Pearson correlation within the matrix sub-compartments was higher (between 0.65-0.79); however, there was a significant difference in the Pearson correlation within the mitochondrial matrix (r=0.79) compared to correlations across mitochondrial matrix and RNP granule (r=0.64, P<0.0001), or compared to correlations across mitochondrial matrix and mitochondrial ribosome (r=0.76, P=0.0198). This analysis indicates that, in general, we do not see major effects due to tagging and bait expression over endogenous levels, validating the Jaccard analysis and demonstrating the predictive value of the dataset.

### Prey-centric analysis identifies functional modules

Prey proteins that are in close proximity to one another should be similarly labeled across all the baits in this dataset, highlighting potential associations in functional complexes or modules. A prey-wise correlation profile should therefore reveal these associations, as was previously done for P-bodies, stress granules and a global map of a human cell (Youn et al., 2018)(https://www.biorxiv.org/content/10.1101/796391v1). We therefore performed a Pearson correlation analysis of the dataset (see details in Methods for parameters and cutoffs) using ProHits-viz (Knight et al., 2017). The resulting heatmap (**Fig. 4A**) of 657 preys enabled us to clearly identify clusters of proteins belonging to either OMM (**Fig. 4B**), IMS/IMM (**Fig. 4C, D**) or matrix sub-compartments (**Fig. 4E**). Analysis of the OMM cluster (**Fig. 4B**) showed the presence of four smaller clusters of proteins (as defined by the hierarchical tree) participating in different biological processes, annotated by the GO terms: organelle organization, nuclear and ER membrane network, establishment of protein localization or cellular response to stress. Detailed analysis of the cluster “organelle organization” (**Supplementary Fig. S4A)** showed a clear clustering of OMM proteins implicated in mitochondrial fusion or fission (**Fig. 4F**). Two other clusters of OMM proteins were also detected: one contained proteins annotated as belonging to several cellular compartments (**Fig. 5A**, see details below) and the other involved in response to stress (**Supplementary Fig. S4B**).

**Figure 4.**
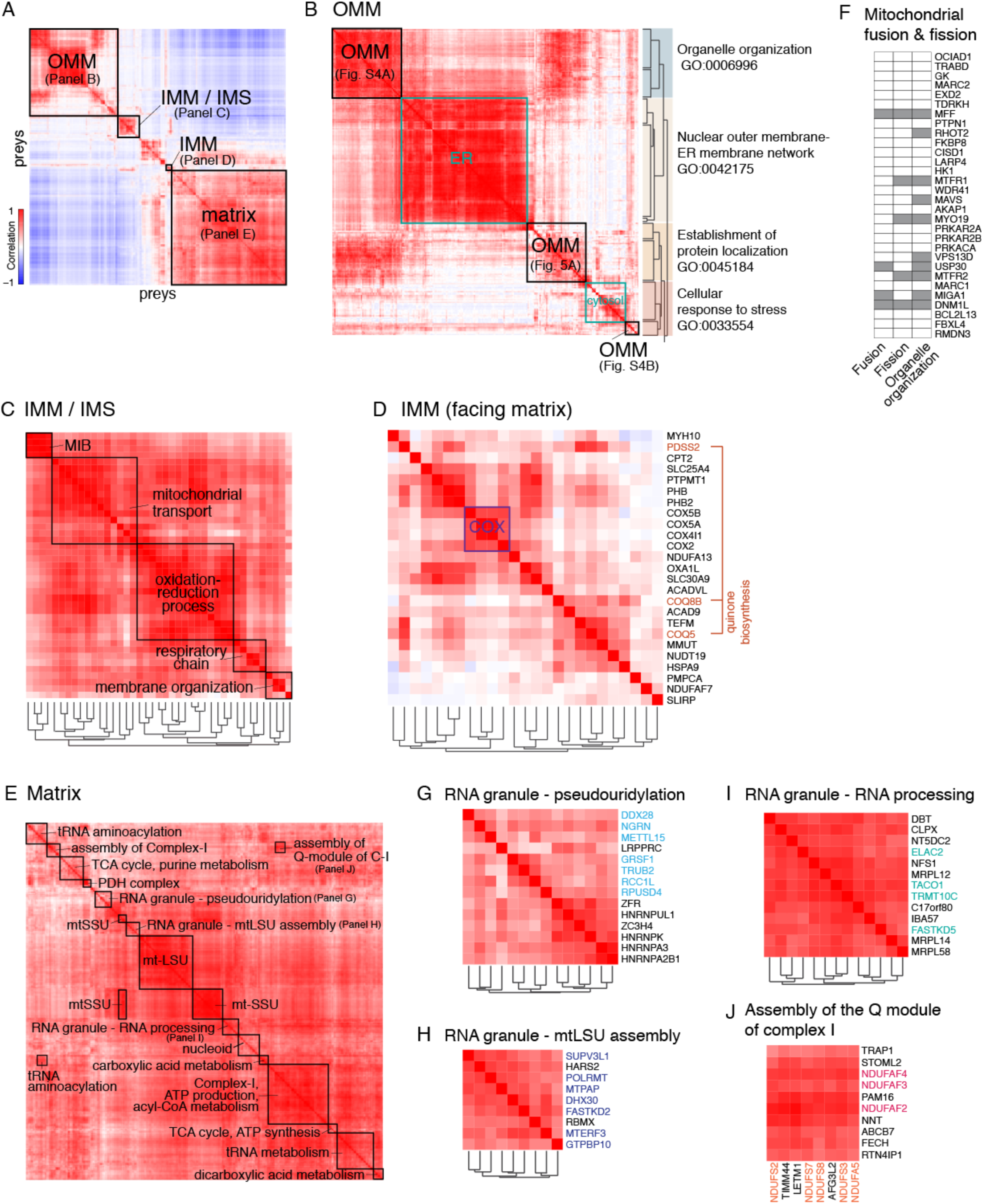
Prey-prey correlation analysis. **A**. A heatmap of the entire network. Discrete clusters, corresponding to mitochondrial outer membrane (OMM), mitochondrial intermembrane space/inner mitochondrial membrane (IMS/IMM), IMM and the matrix are indicated. **B-E**. Enlarged views of OMM (**B**), IMS/IMM (**C**), IMM (**D**) and matrix (**E**) with indicated smaller functional clusters. (**B**) Hierarchical tree and the annotated Panther GO terms Biological process for the four major clusters are indicated to the right of the heatmap. (**C**) Clusters of proteins annotated to be involved in specified functions by Panther GO term analysis. MIB; mitochondrial intermembrane space bridging complex. (**D**) Cytochrome *c* oxidase (COX) cluster is indicated in purple, and proteins involved in quinone biosynthesis in brown. (**E**) Functional clusters annotated according Panther GO term analysis. (**F**). Preys from the cluster “Mitochondrial fusion and fission” (**Supplementary Fig. 4A**) were annotated according to their role in mitochondrial fusion, mitochondrial fission or organellar organization (in grey). **G-J**. Functional clusters. (**G-I**) RNA granule clusters and their hierarchical organization. Proteins previously known to localize to RNA granules are indicated in blue, purple or green, respectively. (**J**) Off-diagonal cluster of preys showing a functional correlation between structural subunits of complex I belonging to the Q-module (shown in orange) and the known Q-module assembly factors (shown in red).

**Figure 5.**
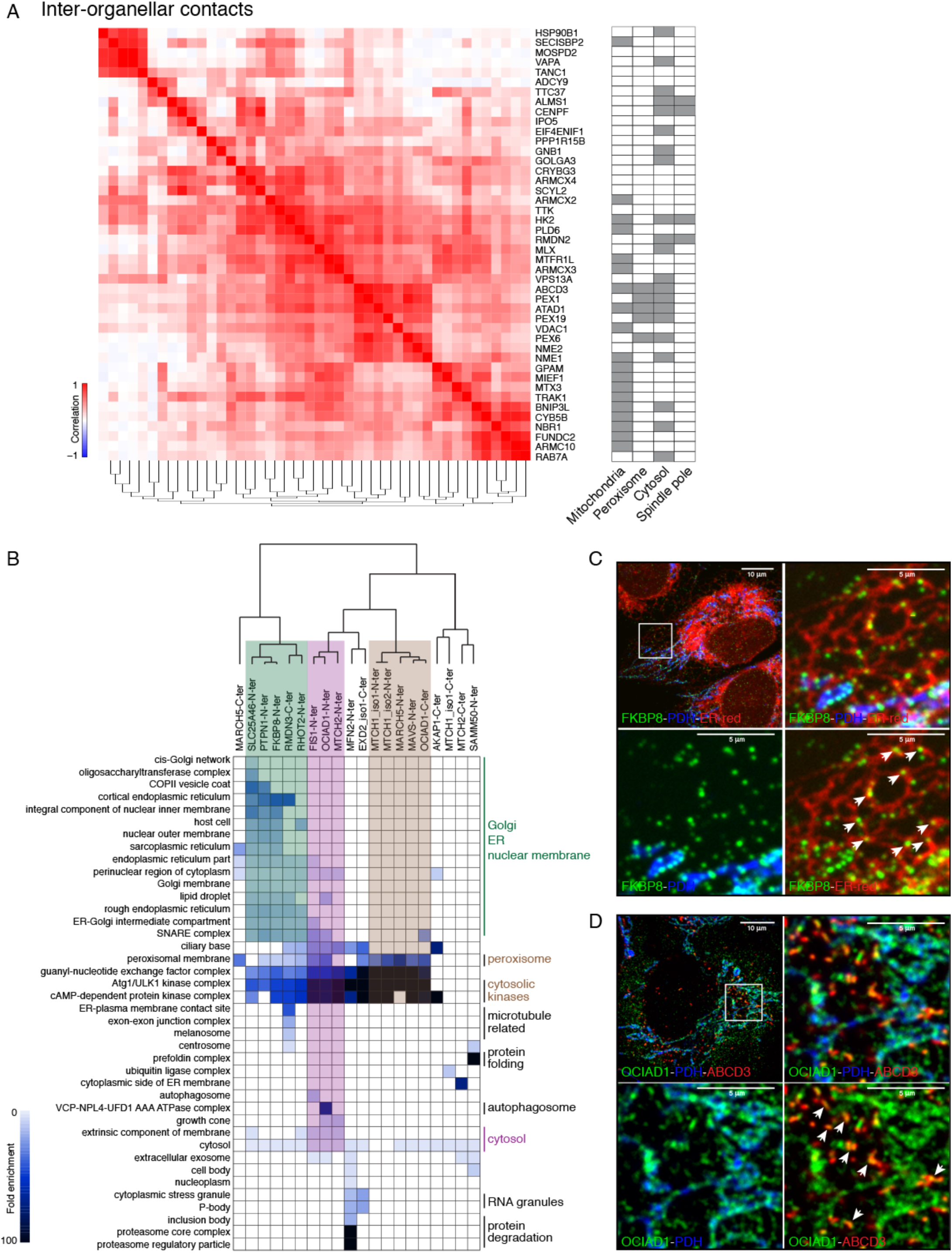
Interaction of mitochondria and other organelles. **A.** A close-up of the OMM prey-prey correlation cluster annotated as “Establishment of protein localization”. Individual proteins were profiled for their cellular localization, which are denoted on the right (in grey). **B.** Heatmap representation of the enrichment values for indicated cellular compartment GO terms for non-Mitocarta preys across mitochondrial outer membrane baits. Bait cluster analysis (dendogram) shows a clear partitioning of individual OMM baits. A cluster of baits playing a role in mitochondrial-ER contacts is highlighted by a green box, proteins involved in mitochondria and peroxisomes are highlighted by a brown box, and proteins interacting predominantly with soluble cytosolic proteins are highlighted in magenta. **C, D**. Dual localization of the FKBP8 (**C**) and OCIAD1 (**D**). Immunocytochemistry was performed on human fibroblasts using antibodies against the endogenous proteins. Representative images of the whole cell (top left panel) with the indicated area used for the zoomed image (top right and bottom panels). (**C**) FKBP8 localizes to both mitochondria (bottom left panel) and ER (white arrows, bottom right panel). ER was visualized using ER-targeted-mCherry constitutively expressed in the human fibroblast line. (**D**) OCIAD1 co-localizes with mitochondria (bottom left panel) and peroxisomes (white arrows, bottom right panel). PDH and ABCD3 were used as a mitochondrial and peroxisomal markers, respectively.

Prey-prey analysis identified two clusters of IMM proteins (**Fig. 4A**), one containing proteins of IMM and IMS (**Fig. 4C**), that was more closely correlated with the OMM proteins, and the other cluster of IMM proteins facing matrix (**Fig. 4D**), clustering with several matrix proteins. The IMM/IMS (**Fig. 4C**) sub-compartment could be divided into five smaller functional modules, encompassing proteins involved in cristae organization, respiratory electron transport chain, oxidation-reduction processes and transport of molecules (such as calcium) across the IMM (**Supplementary Fig. S4C**), the main functions of proteins residing in IMM or IMS. Analysis of the IMM cluster facing matrix (**Fig. 4D**) showed a high correlation cluster of subunits of cytochrome *c* oxidase complex (COX).

Individual clusters within the matrix are indicated in **Fig. 4E**, and encompass major metabolic processes known to occur in the matrix. In particular, we see three clusters of proteins that localize to mitochondrial RNA granules (**Fig. 4G-I**). The mitochondrial RNA granule, a non-membrane delimited structure juxtaposed to the mitochondrial nucleoid, serves as a hub for posttranscriptional RNA processing and modification, and a platform for mitochondrial ribosome assembly (Antonicka and Shoubridge, 2015). Three RNA granule clusters identified in our prey-prey correlation analysis can be distinguished by their functional associations: pseudouridylation (**Fig. 4G**), mtLSU assembly (**Fig. 4H**) and RNA processing (**Fig. 4I**); however, all three RNA granule modules correlate highly with proteins of the mitochondrial ribosome. A high specificity of interaction (enrichment) among the pseudouridine synthase module proteins (Antonicka et al., 2017) can be also visualized in a specificity plot, as shown for RCC1L (WBSCR16) (**Supplementary Fig. 4D**), a protein of yet undefined function within the RNA granule. Interestingly, this plot shows a single proximity interactor with infinite specificity, NME6, a dinucleotide kinase of unknown function that was previously shown to interact with RCC1L by affinity enrichment-mass spectrometry (Floyd et al., 2016).

Analysis of the prey-prey heatmap revealed the presence of off-diagonal clusters, indicating that proteins belonging primarily to two diverse groups, due to, for example, their localization within mitochondria, or within a macromolecular complex, have similarities in their labeling profiles (**Fig. 4J, Supplemental Fig. S4E**). **Fig. 4J** shows one of the off-diagonal clusters identified in the matrix sub-compartment: Complex I subunits (NDUFA5, NDUFS2, NDUFS3, NDUFS7 and NDUFS8) which form the so-called Q-module of Complex I (Guerrero-Castillo et al., 2017), distinctly correlate with the Q-module assembly factors (NDUFAF2, NDUFAF3, NDUFAF4). These data indicate that prey-prey analysis is suitable for assigning proteins to different spatial or functional networks, as well as for detection of putative associations between such networks. The ability of this unsupervised correlation analysis to accurately cluster known protein complexes as well as functionally related proteins validates the robustness of our dataset for practical use in assigning proteins to their proximal environment or functional network.

### OMM baits identify specific interactions with other organelles

Mitochondrial OMM baits that faced the cytosol detected not only mitochondrial preys, but also proximity interactors belonging to other cellular compartments. The median number of non-mitochondrial preys detected per OMM bait was 42 (**Fig. 2B**). The prey-prey correlation heatmap (**Fig. 4B**) showed a cluster of preys that were annotated by the GO terms Establishment of protein localization (GO:0045184) and Transport (GO:0006810). Several preys in this cluster were assigned to more than one organelle or as belonging to more than one cellular compartment (**Fig. 5A**). This suggests that these proteins either localize to multiple compartments (possibly as different isoforms), that they are involved in inter-organellar contact sites, or transport between organelles.

To determine which organelles the OMM baits interacted with, we proceeded to analyse all identified non-mitochondrial preys for each OMM bait for their GO terms: Cellular Compartment (PANTHER), and an enrichment score was assigned to each GO term based on the fold enrichment score in PANTHER. The most specific subclass terms were used for further analysis (**Supplementary Table S6**). The resulting heatmap (**Fig. 5B**) between OMM baits and preys matched GO terms shows clear clustering of individual baits based on the organelle they predominantly interact with. SLC25A46, PTPN1 (known to be also present in the ER), FKBP8, RMDN3 and RHOT2 detect close-proximity preys primarily from the endoplasmic reticulum, or associated with the nuclear membrane or Golgi, while MTCH1, MARCH5, MAVS and OCIAD1 detected preys from peroxisomes. RMDN3 has been shown to tether mitochondria and ER via its interaction with VAPB (De Vos et al., 2012) and SLC25A46 is necessary for efficient lipid transfer between ER and mitochondria (Janer et al., 2016). MAVS localizes to both mitochondria and peroxisomes and acts as a signalling platform for antiviral innate immunity (Dixit et al., 2010). These examples, which corroborate our correlation analysis, suggest either dual/multiple localization of several OMM proteins, or that the contact sites they form between mitochondria and other organelles are specific to the organelle involved. Further functional analysis should provide some insight into the different roles these contact sites play in the regulation of cellular metabolism.

Immunofluorescence analysis of individual baits in 293 Flp-In T-REx cells suggests that some baits might localize to more than one cellular compartment (**Supplementary Fig. S5A**). We chose two such baits, FKBP8 and OCIAD1, to determine whether they also localize to ER and peroxisomes, respectively, the compartments in which their detected preys were enriched. Using antibodies against endogenous FKBP8 and OCIAD1, together with markers for mitochondria, ER and peroxisomes, we were able to show that FKBP8 localized on the periphery of both mitochondria and ER (**Fig. 5C**). OCIAD1 was predominantly localized to mitochondria; however, a fraction of immuno-detectable OCIAD1 co-localized with the peroxisomal marker ABCD3 (**Fig. 5D**), but not an ER marker (**Supplementary Fig. S5C**), consistent with our BioID results.

Taken together, our data show that individual mitochondrial OMM proteins interact with unique subsets of cellular proteins and confirm that BioID can be used to determine the identity of proteins on mitochondria that form contact sites with other organelles. The five baits that predominantly recognized ER preys (SLC25A46, PTPN1, FKBP8, RMND3, RHOT2), each identified between 182-230 preys, 72 of which were in common (**Supplementary Table S6**), and represent proteins that might be enriched at ER-mitochondrial contact sites. More than half of these 72 preys overlapped with ER-mitochondria contact proteins reported in one or more recent studies (Hung et al., 2017), Cho et al. (https://doi.org/10.1101/2020.03.11.988022)(Kwak, 2020) (**Supplementary Figure. S5D, Supplementary Table 6**). The remaining 32 proteins, which we designate as BioID mitochondria-ER orphans, provide a valuable source of candidates for further functional validation.

### Sub-compartmental localization of EXD2 isoforms in mitochondria

While bait-bait cluster analysis (**Fig. 3**) showed a clear separation between baits of the mitochondrial matrix and the OMM, surprisingly, one of the OMM baits, EXD2, showed a partial correlation with matrix baits, predominately structural subunits of the mitochondrial ribosome (**Fig. 6A, C**). A possible explanation for this behavior could be that this protein is dually localized in different mitochondrial sub-compartments, an extremely rare occurrence. EXD2, Exonuclease 3’-5’ Domain Containing 2, is a protein that has been described to localize to the OMM (Hensen et al., 2018; Park et al., 2019), the mitochondrial matrix where it was described to play a role in mitochondrial translation through its RNA endonuclease activity (Silva et al., 2018), as well as the nucleus where it was suggested to play a role in double-strand break resection and (Broderick et al., 2016; Hensen et al., 2018; Nieminuszczy et al., 2019; Park et al., 2019; Silva et al., 2018). Two EXD2 isoforms are predicted in databases (**Fig. 6B**), resulting from an alternative splicing (skipping of exon 2), producing an isoform 2 lacking the first 125 amino acids. An identical protein would also arise if a downstream methionine, M126, were used for translation initiation, and such alternative translation initiation could explain our BioID data (as the tagged bait is not spliced). The second isoform lacks a large part of the 3’-5’ endonuclease which extends from aa 76-295, but still contains the HNH endonuclease domain from aa 408-470 (Park et al., 2019). The presence of several EXD2 isoforms (or products from posttranslational processing) was confirmed using stealth siRNA to knock-down EXD2 in cells (**Supplementary Fig. S6A**). To test whether alternative translation initiation from M126 of EXD2 isoform 1 could be responsible for the proximity-labelling of matrix baits, we substituted alanine for methionine at position 126 and created three BirA*-tagged constructs: wild-type isoform 1 (iso1), isoform 1 carrying the M126A variant (M126A), and wild-type isoform 2 (iso2). All three proteins localized to mitochondria (**Supplementary Fig. S6B, Fig. 6B**). A proteinase K protection assay confirmed that iso1 and M126A localized to the outer mitochondrial membrane, while isoform 2 localized inside of mitochondria (**Fig. 6B**). Iso1 and M126A contain a predicted transmembrane domain between amino acids 7 and 25, not present in iso2, and we confirmed that both iso1 and M126A were integral membrane proteins, while iso2 was predominantly soluble (**Supplementary Fig. S6C**).

**Figure 6.**
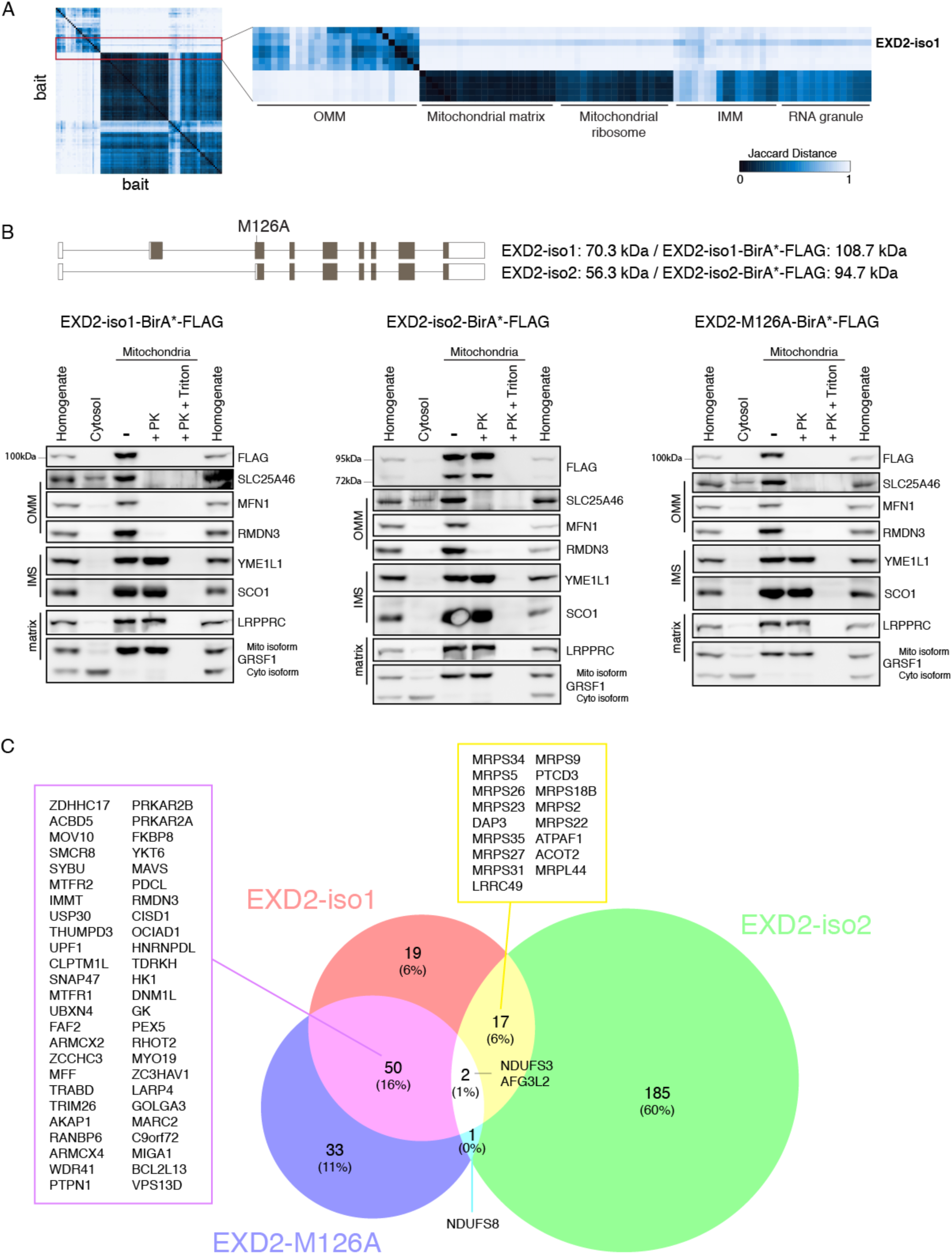
Determination of the sub-compartmental localization of EXD2-BirA* isoforms. **A.** Zoom of the indicated area of the bait-bait heatmap (same figure as in Fig. 3) shows a correlation between OMM bait EXD2-iso1 and mitochondrial matrix baits. The Jaccard distance score between EXD2-iso1 and matrix baits was between 0.75 and 0.9, while the Jaccard distance score between all other OMM baits and matrix baits was between 0.9 and 1. **B.** The sub-mitochondrial localization of EXD2-BirA*-FLAG-tagged isoforms was established using a proteinase K protection assay of isolated mitochondria from Flp-In T-REx 293 cells stably expressing EXD2-BirA*-FLAG-tagged isoforms. FLAG antibody was used for visualization of EXD2 isoforms, and antibodies against indicated proteins from individual sub-mitochondrial compartments were used as markers. Schematic presentation of the EXD2 isoforms with specified sizes for the predicted proteins. **C.** Venn diagram of the BioID results for all three EXD2 isoforms confirms the mitochondrial localization of the isoforms. The list of preys common to at least two isoforms is shown, validating the OMM localization of EXD2 iso1 and M126A, and a partial overlap of detected preys between EXD2 iso1 and iso2.

BioID analysis (**Fig. 6C, Supplementary Table S7**) confirmed that iso1 and M126A localize in OMM, while iso2 localizes to the matrix. While 19 proteins were common to iso1 and iso2, all from the IMM or matrix, the spectral counts detected for these proteins were generally 4-8 times lower for iso1 than iso2 (**Supplementary Fig. S6D**), indicating that translation from the downstream ATG is much less efficient than translation of the full-length isoform. The M126A construct did not identify any matrix preys, validating our prediction that alternate start site at position M126 is indeed responsible for the identification of matrix preys with iso1. The loss of the interaction with matrix proteins by M126A was not due to a decreased expression of the bait (**Fig. 6B**); as both iso1 and M126A baits detected a similar number of preys with a BFDR*≤*1% (88 preys for iso1; 86 preys for M126A) and 60% of detected preys were common to both baits (**Fig. 6C**). The common preys between these two baits were mostly preys of the OMM or assigned to a different cellular compartment, as the BirA*-tag faced the cytosol (peroxisomal membrane, cytosolic kinases, cytoplasmic stress granule); however, no nuclear preys were detected, confirming the exclusive mitochondrial localization of EXD2. Only two IMM preys were common to all three baits, AFG3L2 and NDUFS3; however, the number of detected spectral counts by iso1 and M126A was 10-times lower than for iso2 (**Supplementary Table S7**). These results reinforce the strength of our analysis and dataset, in that we are able to see a clear difference between individual baits in their localization and the proximity partners they detect using BioID.

### Mitochondrial ribosome proximity network and ribosome assembly

Prey-prey correlation analysis of detected preys using Fold Change as the abundance measure detected 78 mitochondrial ribosomal proteins forming individual clusters within the mitochondrial matrix (**Fig. 7A, Supplementary Fig. S7**). MRPS36 protein, which was not detected as a subunit of the human mitochondrial ribosome in cryo-EM analyses (Amunts et al., 2015), has rather been shown to be part of the alpha-ketoglutarate complex (Heublein et al., 2014), and it clustered in our analysis with OGDH, the E1 component of the alpha-ketoglutarate complex, validating previously published results. A Cytoscape view of the correlation matrix revealed a network of mitochondrial ribosomal proteins and their correlated partners (**Fig. 7B**), where close relationship among the proteins of small ribosomal subunit (mtSSU) were readily observed. Proteins of the large ribosomal subunit (mtLSU) formed three largely discrete clusters based on the prey-prey correlation analysis (**Fig. 7A, Supplementary Fig. S7)**. We displayed these clusters of proteins on the cryo-EM structure of the human mitochondrial ribosome (PDB 3J9M, mtLSU shown only), and their location within the structure are compatible with a potential modular assembly of the mtLSU. The cluster L3 (**Fig. 7B, Supplementary Fig. S7B**) shows a close correlation between MRPL11/MRPL54 and MALSU1, previously shown to be an assembly factor of the mitochondrial large ribosomal subunit (Fung et al., 2013; Rorbach et al., 2012) and suggests a possible role of MALSU1 in the addition of MRPL11/MRPL54 to the ribosome. Several other factors involved in ribosome assembly were also identified in the cluster (RPUSD4, GTPBP10, DDX28, GRSF1). The only mtLSU protein not identified in this analysis was MRPL36, which was not detected as a prey in our or other publicly available BioID analyses. Recently, a low-resolution model of mitoribosome assembly was suggested using pulse-chase SILAC (Bogenhagen et al., 2018), categorizing proteins into early, intermediate and late proteins for mtLSU assembly. Our data correlate well with the suggested model, as the mtLSU-cluster 1 contained early and intermediate proteins, the mtLSU-cluster 2 contained early and late proteins, and the mtLSU-cluster 3 contained late proteins.

**Figure 7.**
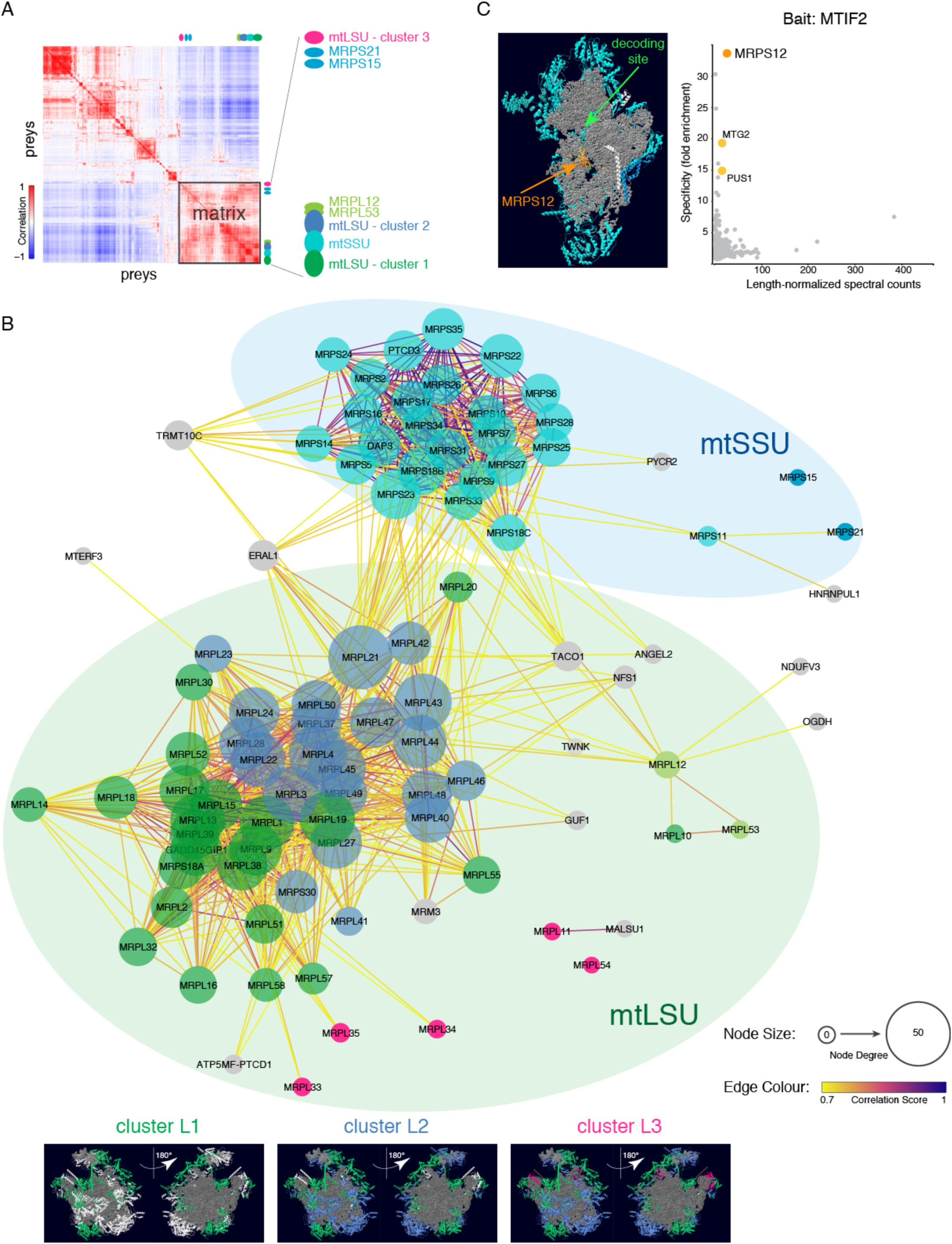
Mitochondrial ribosome interaction network. **A.** A prey-prey heatmap based on Fold Change enrichment revealed 78 mitochondrial ribosomal proteins forming high-correlation clusters within the matrix cluster (**Supplementary Fig. S7**). Proteins of the small ribosomal subunit (mtSSU) are indicated in turquoise and light blue, proteins of the large ribosomal subunit (mtLSU) are in dark blue, green and pink. **B.** A mitochondrial ribosome network diagram was created using Cytoscape based on a correlation matrix from the prey-prey analysis. Only correlations greater than 0.7 were considered for the network, and only edges with at least one mitochondrial ribosomal protein were included. The edges are colour-coded based on the correlation score. Node size shows the degree of connectivity of each node, and is defined as a number of edges linked to a node. mtSSU proteins are indicated in turquoise and light blue; and the mtLSU proteins are in dark blue, green and pink. Other interactors with a correlation greater than 0.7, which are not part of mitochondrial ribosome, are indicated in grey. Proteins of the mtLSU identified in individual clusters were displayed on the cryo-EM structure of the human mitochondrial ribosome (PDB 3J9M, mtLSU shown only). The solvent side of the ribosome is shown on the left, and the interphase side is shown on the right. The 16S rRNA and mtRNA-Val are depicted in grey. **C.** Interphase view of the mtSSU with proteins detected in (**A**) depicted in turquoise and light blue. The position of MRPS12 and the decoding site are also indicated. The 12S rRNA is shown in grey. Specificity plot of the MTIF2 bait shows a highly specific interaction with MRPS12. Two other proteins involved in mitochondrial translation with a specificity greater than ten, MTG2 and PUS1, are also indicated (in yellow), all other detected prey are in grey.

The mtSSU proteins largely clustered together with the exception of MRPS15 and MRPS21 (**Fig. 7A and 7B, Supplementary Fig. S7A**), both of which were previously described as late assembly partners for mtSSU (Bogenhagen et al., 2018). Three proteins of the mtSSU (MRPS12, MRPS37 and MRPS38) were not identified in this analysis. MRPS37 (CHCHD1) and MRPS38 (AURKAIP1) were not detected as preys in our or other published BioID analyses. Interestingly, MRPS12, a protein present at the interphase side of the mtSSU in the vicinity of the mRNA decoding site (**Fig. 7C**), was detected by BioID analysis as a specific interactor of the mitochondrial initiation factor MTIF2 (it was not present in the prey-prey correlation heat map, as the analysis required a minimum of two baits detecting a prey for the prey to be included in the correlation matrix). A reciprocal interaction between MRPS12 and MTIF2 was also detected, when MRPS12 was used as a bait. MTIF2 promotes the binding of initiator tRNA to the mtSSU (Liao and Spremulli, 1991). These data suggest that MRPS12 might be responsible for bringing MTIF2 together with the initiator tRNA, and possibly mitochondrial mRNAs to the active site of the ribosome for translation initiation, indicating that our dataset provides a valuable input into a possible mechanistic route of mitochondrial translation initiation, a process whose regulation remains poorly understood in mammalian mitochondria.

## Discussion

In this paper we describe the creation of mitochondrial proximity interaction network using 100 individual baits from all mitochondrial compartments and show several examples in which we apply our BioID data. Using less than 10% of the mitochondrial proteome as baits, we were able to identify more than half of the proteins annotated in Mitocarta 2.0 as high-quality proximity interactors. Defining sub-organellar environments (matrix, IMS) allowed us to investigate specific proximal interactors for individual baits, providing potential insight into functionality for proteins, or those with poorly characterized functions. For instance, the mitochondrial translation termination factor MTRF1L not only enriched for two subunits of the mitoribosome in the structure responsible for recruiting mitochondrial translation factors, (MRPL10, MRPL53) and a GTPase involved in mitoribosome assembly (GTPBP10), but also three complex I assembly factors (**Fig. 1C**). One of the complex I assembly factors, ACAD9, enriched for other assembly complex I factors in the same module, and interestingly three subunits of PDK (pyruvate dehydrogenase kinase), the kinase responsible for phosphorylating and inactivating PDH (pyruvate dehydrogenase), slowing entry of pyruvate into the TCA cycle. These data suggest a close functional coupling between mitochondrial translation, assembly of the first complex in the OXPHOS system, and control of the generation of NADH, by regulating activity of the TCA cycle, a hypothesis that deserves further functional testing. While other proximity labelling techniques, like APEX, are very useful tools for cataloguing the matrix and IMS proteome (Hung et al., 2014; Rhee et al., 2013), they do not permit the same kind of specificity analysis as BioID, which can serve to identify potential interactors for downstream functional analyses.

Bait-bait analysis showed that we have the resolution to assign protein complexes or functional protein modules to sub-compartments within mitochondria. This analysis very clearly separated OMM, matrix and IMM compartments, and showed clustering of sub-compartments in the matrix space associated with mitochondrial translation, mitoribosome assembly, and the RNA granule. We have previously investigated the mitochondrial RNA granule, a non-membrane delimited structure that houses much of the enzymatic machinery for posttranscriptional processing of the polycistronic RNAs transcribed from the mitochondrial genome, and for mitochondrial ribosome biogenesis. Bait-bait analysis clustered together some of the main protein components of the RNA granules including GRSF1, DDX28, DHX30, FASTKD2, that we previously identified by immunoprecipitating endogenous GRSF1 after UV-crosslinking, a completely orthogonal technique (Antonicka and Shoubridge, 2015), while other proteins that were also identified in the granule such as NGRN, RCC1L, FASTKD5, MTERFD1, and RPUSD4 clustered with the mitochondrial ribosome, suggesting that the RNA granule may be further structured, an idea that could be tested by combining BioID with XL-MS.

A comprehensive analysis of the preys identified by all 100 baits allowed us to identify some potentially novel mitochondrial proteins (some of which were confirmed while this study was in progress). We also identified a few proteins in the IMS and matrix that clearly had multiple cellular locations that could not be readily visualized by immunofluorescent analyses, possibly because of skewing of their distribution, a hypothesis that would require further testing. GRSF1, an RNA-binding protein, which appears exclusively mitochondrial by immunofluorescence, showed clear proximity interactors in other compartments including the nucleolus. Such dual or multiple cellular localizations could result from alternate, downstream ATG start sites, or alternatively from multiple protein isoforms. Another example of dual targeting, but within the organelle itself, emerged from the analysis of the proximity interactome of EXD2. This protein, which has both exo- and endonuclease domains, had previously been proposed to localize to the nucleus, the OMM or the matrix, with different functional attributions. Our data clearly show that while EXD2-iso1-BirA* is predominately an OMM protein, a fraction localizes to the matrix, likely representing an isoform that is translated from an alternate downstream ATG start site. These data suggest that the precise function of EXD2 deserves further investigation.

A particularly informative analysis of our data comes from prey-prey clustering. This correlation analysis predicts a clustering of preys involved in the same pathways or modules, and we validate this for proteins involved in mitochondrial fission and fusion, assembly of the respiratory chain complexes, the nucleoid, the RNA granule amongst others. While most of the baits in the matrix, IMM, and IMS identified preys within the organelle, as expected most OMM baits identified preys in the cytosol or proteins associated with other organelles. Using GO terms we identified clusters of proteins known to be involved with particular biological processes, but these clusters often included uncharacterized proteins whose biological functions could be investigated by detailed functional analysis. Specificity plots are particularly useful in identifying preys that are specifically enriched for particular baits, and can help in identifying novel proteins that associate with particular modules as we show with the pseudouridylation module (**Supplementary Fig. S4D**) and for several other proteins (**Fig. 2D, Supplementary Fig. S2C**). These data represent a rich source from which to generate mechanistic hypotheses regarding the role of a large number of poorly characterized proteins associated with mitochondrial metabolism and morphology.

Although we choose very few structural subunits of the mitochondrial ribosome as baits, we identified nearly all of them as preys, mapped these onto a cryo-EM structure of the ribosome, and remarkably showed that they cluster in the prey-prey analysis in a manner that is consistent with a model for its assembly. The clusters also identified a number of known ribosome assembly factors, and uncharacterized proteins that may also be involved in this process.

What was perhaps not expected for the prey-prey analysis of OMM baits is the specialization of proximity interactions. Unsupervised clustering of detected non-Mitocarta preys across the OMM baits showed clear separation of OMM proteins, with some interacting with the ER and associated endo-membranes, while others showed clear enrichment with peroxisome markers. Others, such as MFN2, which has been proposed to act as a mitochondrial-ER tether (de Brito and Scorrano, 2008), showed a strong enrichment with the proteosomal machinery, and no interaction with ER proteins, suggesting that its role as a mitochondrial-ER tether might deserve further scrutiny. We were also able to validate dual localization of two OMM proteins to ER (FKBP8) or peroxisomes (OCIAD1).

The area of organelle contact sites has become an area of intense investigation as these sites are important platforms for lipid exchange calcium movement, especially between mitochondria and the ER (Csordas et al., 2018). Our data suggest that mitochondria-organelle contact sites may in fact be exquisitely tailored to specific cellular functions with unique proteomes that depend on the specific nature of the contact, be it with the ER or peroxisomes. BioID could potentially be used to profile alterations in the proximity interactomes at contact sites in response to physiological challenges or mutations in genes coding for proteins that have a role in organelle tethering or metabolite exchange.

Among the 32 BioID mitochondria-ER orphans we identified (**Supplemental Fig. S5D**) are three members of the SMCR8-C9orf72-WDR41 complex, which has been implicated in autophagosome maturation (Yang et al., 2016), selective autophagy (Sellier et al., 2016), and both positive and negative regulation of autophagy (Jung et al., 2017; Ugolino et al., 2016). The transfer of calcium from the ER to mitochondria, which occurs at contact sites, regulates autophagosome formation (Gomez-Suaga et al., 2017; Mallilankaraman et al., 2012; Vicencio et al., 2009; Wong et al., 2013), and autophagosomes form at sites of apposition between ER and mitochondria (Garofalo et al., 2016; Hailey et al., 2010; Hamasaki et al., 2013), all consistent with the presence of the SMCR8-C9orf72-WDR41 complex at mitochondria-ER contact sites.

We also identified PRKAR2A and PRKAR2B, the regulatory subunits of the cAMP-dependent protein kinase complex (PKA) as mitochondria-ER orphans. The outer mitochondrial membrane harbours cAMP/PKA signalling microdomains consisting of the A-kinase anchoring protein AKAP1, which serves as a scaffold to recruit PKA (Feliciello et al., 2005; Lefkimmiatis and Zaccolo, 2014). PKA activity regulates several mitochondrial processes (Ould Amer and Hebert-Chatelain, 2018), and of particular interest mitochondrial fission and fusion (Flippo and Strack, 2017). Mitochondrial fission requires the recruitment of the GTPase DNM1L by its adaptator MFF at mitochondria-ER contact sites (Osellame et al., 2016). PKA phosphorylates DNM1L and inhibits its capacity to induce mitochondria fission, and thus serves as a mechanism to regulate the balance between fission and fusion (Cribbs and Strack, 2007). Our analyses also identified AKAP1, DNM1L and MFF as enriched mitochondria-ER contact proteins (**Supplementary Table S6)**, all of which were seen in at least one other study profiling proteins at contact sites (Hung et al., 2017)(Cho et al. (https://doi.org/10.1101/2020.03.11.988022) (Kwak, 2020). Thus, our BioID analysis of proximity interactors at ER-mitochondria contacts is able to identify much of the molecular machinery that integrates cAMP signalling to regulate mitochondrial fission-fusion events.

As all data are available to the readers, our dataset provides an enormous source of information and can be used directly (to find proximity interactors of the protein of interest, if in the dataset) or indirectly (to compare reader’s data to our dataset). For easy access to the data, we deposited all data to ProHits (http://prohits-web.lunenfeld.ca), where information on individual baits and their detected preys can be found.

## Acknowledgments

We thank Kathleen Daigneault for the technical work in this project (generation of most of the BioID constructs and cell lines), Mari Aaltonen for critical reading of the manuscript and the members of Shoubridge lab for helpful discussions. This research was funded by grants from the United Mitochondrial Disease Foundation and the Canadian Institutes Health Research to EAS. ACG was supported by a Canadian Institutes of Health Research (CIHR) Foundation Grant (FDN 143301). Proteomics work was performed at the Network Biology Collaborative Centre at the Lunenfeld-Tanenbaum Research Institute, a facility supported by Canada Foundation for Innovation funding, by the Ontarian Government and by Genome Canada and Ontario Genomics (OGI-097, OGI-139). ACG is the Canada Research Chair in Functional Proteomics.

## Author Contributions

H.A., E.A.S. and A.-C.G. conceived the project. H.A., E.A.S. and A.-C.G. wrote the paper.

H.A. supervised preparation of all BioID samples.

Z.-Y.L. performed affinity purification and mass spectrometric acquisition.

H.A., A.J. and W.W. performed Western blot and immunofluorescence studies and analyzed the data.

H.A. and A.-C.G. performed data analysis.

E.A.S. and A.-C.G. supervised the project.

## Figure legends

**Supplementary Figure S1** (related to Fig. 1)

**A.** Immunofluorescence images of mitochondrial targeting of MTS-BirA* constructs. FLAG staining in green, mitochondrial marker TOMM20 in red, DAPI in blue. Scale bar, 10 μm.

**B.** Specificity of coverage of the Matrix-BirA* compared to Matrix-APEX (Rhee et al., 2013). Preys detected by Matrix-BirA* were compared to either Mitocarta 2.0 or GOCC to determine mitochondrial specificity. 249 preys (93%) were previously annotated as mitochondrial proteins. Matrix specificity was calculated as described in (Rhee et al., 2013): Matrix-BirA* and MTS-APEX were compared to a dataset with 424 proteins with sub-mitochondrial localization. Sub-mitochondrial annotation was available for 142 Matrix-BirA* detected preys, resulting in 99% matrix specificity.

**C.** Pairwise comparison of high-confidence identified preys by MTS-COX4 vs. MTS-COX8 (**left**) and by MTS-OTC vs. MTS-COX4 (**right**) to identify the common and specific interactors for the baits (log_2_ transformed average spectral counts normalized to total abundance). The pairwise correlation coefficient, calculated from the untransformed spectral count matrix of the two conditions, is indicated. Proteins annotated by a GO term “mitochondrial translation elongation” (GO:0070125) are indicated in light blue, “mitochondrial genome maintenance” (GO:0000002) in purple and “RNA modification” (GO:0009451) in green, and selected significantly different interactors belonging to these groups are indicated by their gene name. Preys in these groups which passed BFDR for both baits are outlined with a full circle black border, proteins which passed BFDR only for one bait are indicated by a half circle border. Interactors in beige represent preys where no spectra were detected for the other bait, while interactors in light grey passed the BFDR*≤*1% only for one bait. Interactors common to both baits are in dark grey.

**D.** Specificity of coverage of the IMS-BirA* compared to IMS-APEX (Hung et al., 2014). Preys detected by IMS-BirA* baits were compared to either Mitocarta 2.0 or GOCC to determine mitochondrial specificity. 112 preys (93%) were previously annotated as mitochondrial proteins. Potential “IMS exposure” was calculated as described in (Hung et al., 2014): IMS-BirA* and IMS-APEX were compared to a dataset with 579 proteins with sub-mitochondrial localization. Sub-mitochondrial annotation was available for 68 IMS-BirA* detected preys, resulting in 97% “IMS-exposure”.

**E.** Characterization of the mitochondrial IMS “environment”. Venn diagram of two IMS-BirA* baits showing the most common set of interactors, the “environment”. Top 40 preys (based on normalized spectral counts) are indicated. The 5 preys in bold are common to both the matrix and IMS top 40 list; and are IMM proteins.

**F.** Pairwise comparison of high-confidence identified preys by MTS-OPA1 vs. MTS-AIFM1 to identify common and specific interactors for the baits (log_2_ transformed average spectral counts normalized to total abundance). The pairwise correlation coefficient, calculated from the untransformed spectral count matrix of the two conditions, is indicated. Proteins annotated by a GO term “protein targeting to mitochondrion” (GO:0006626) are indicated in orange, and significantly different interactors belonging to this groups are indicated by their gene name. Preys in this group which passed BFDR for both baits are outlined with a full circle black border, proteins which passed BFDR only for one bait are indicated by a half circle border. The baits are in blue. Interactors in beige represent preys where no spectra were detected for the other bait, while interactors in light grey passed the BFDR*≤*1% only for one bait. Interactors common to both baits are in dark grey.

**Supplementary Figure S2** (related to Fig. 2)

**A.** Correlation analysis of the total number of identified preys (grey triangles), Mitocarta preys (blue circles) and non-Mitocarta preys (green squares) by individual baits and the expression level of the bait (BirA* spectra). The baits were divided based on their localization within mitochondria. The expression level of the bait was defined as the average of detected spectra for the BirA* tag, as BirA* biotinylates itself. The table indicates the calculated Pearson correlation scores, and corresponding P values. n.s., not significant

**B.** GO term analysis of non-Mitocarta detected preys for two matrix baits (GRSF1 and CHCHD1) and one IMS bait (HAX1). 139 non-Mitocarta preys were detected for GRSF1, 110 preys for CHCHD1, 97 preys for HAX1 and the pie-charts represent the relative distribution of the detected non-Mitocarta preys within the individual GO terms.

**C.** Specificity plot examples of two IMM baits (CCDC90B, SLC25A12-N-ter), one OMM bait (MTCH1_iso2-C-ter) and one matrix bait (ACAD9) indicating the specific interaction with preys identified as single interactors in our dataset. Infinite specificity signifies that no spectrum was detected for that prey in any other sample analyzed.

**Supplementary Figure S3** (related to Fig. 3)

**A**. Pearson correlation analysis between individual baits within identified mitochondrial sub-compartments and within detected clusters as well as between compartments and between clusters. A matrix of the average spectral counts (normalized to total spectral counts per bait) for all high-confidence preys (BFDR*≤*1%) across all baits in the dataset was used to calculate the Pearson correlation matrix for the baits. The mean of Pearson correlations between baits belonging to individual sub-compartments or clusters within these compartments, or between two sub-compartments or clusters are indicated in the figure.

**B-D**. Pairwise comparison of high-confidence identified preys by two matrix baits (**B**), two OMM baits (**C**) and one matrix and one OMM bait (**D)** to identify common and specific interactors for the baits (log_2_ transformed average spectral counts normalized to total abundance). The Pearson correlation coefficient, calculated from the untransformed spectral count matrix of the two conditions, is indicated. Specific interactors in beige represent preys where no spectra were detected for the other bait, while specific interactors in light grey passed the BFDR*≤*1% only for one bait. Interactors common to both baits are in dark grey. Baits are indicated in red.

**Supplementary Figure S4** (related to Fig. 4)

**A, B**. Enlarged views of the functional clusters “Organelle organization” and “Cellular response to stress” with individual proteins indicated on the right and OMM proteins highlighted in blue. Hierarchical tree was used to determine the “Mitochondrial fusion and fission” cluster.

**C.** Proteins from the IMM/IMS cluster (Fig. 4C) annotated to be involved in specified functions by Panther GO term analysis are indicated in grey.

**D.** A specificity plot example of a pseudouridine synthase module protein RCC1L (bait name in our dataset WBSCR16), indicating an enrichment for interaction with other proteins from the pseudouridine synthase module (green) and with other mitochondrial RNA granule proteins (orange). NME6, a mitochondrial protein belonging to a family of nucleoside diphosphate kinases, is an interactor with infinite specificity (no spectrum was detected for this prey in any other sample analyzed).

**E.** Two off-diagonal clusters from the prey-prey analysis (same figure as in Fig. 4A) indicate a close relationship between proteins clustering primarily in IMM/IMS and IMM (bottom) and between clusters of IMM and matrix proteins (right). Individual preys are indicated on the bottom and to the right of the figure. Respiratory chain cluster signifying a close correlation of known cytochrome *c* oxidase (COX) subunits and assembly factors (shown in purple).

**Supplementary Figure S5** (related to Fig. 5)

**A.** Immunofluorescence analysis of three OMM baits, RMDN3-BirA*-FLAG, BirA*-FLAG-FKBP8 and BirA*-FLAG-OCIAD1 in Flp-In T-REx 293 cells. All three baits co-localized with mitochondria, however BirA*-FLAG-FKBP8 and BirA*-FLAG-OCIAD1 show significant staining outside mitochondria. FLAG antibody was used to visualize individual baits and TOMM20 was used as a mitochondrial marker. DAPI, in blue. Scale bar, 10 μm.

**B.** Immunofluorescence analysis of endogenous FKBP8 in human fibroblasts. FKBP8 does not co-localize with the peroxisomal marker ABCD3. PDH, mitochondrial marker. Scale bar, 10 μm.

**C.** Immunofluorescence analysis of endogenous OCIAD1 in human fibroblasts. OCIAD1 does not co-localize with the ER marker, ER-targeted-mCherry. Cytc, mitochondrial marker. Scale bar, 10 μm.

**D.** Identification of mitochondria-ER contact site orphans. Venn diagram comparing our BioID (Mito-ER) preys and previously published proteomic datasets of mitochondria-ER contact site proteins (Hung et al., 2017)(https://doi.org/10.1101/2020.03.11.988022)(Kwak, 2020). BioID (Mito-ER) represents 72 preys common to the cluster of five OMM baits (SLC25A46, PTPN1, FKBP8, RMDN3 and RHOT2) (Fig. 5B, in green), which proximity-label predominantly ER proteins. The numbers in parenthesis represent the total number of proteins in each dataset (**Supplementary Table S6**). The four preys common to all datasets (medium grey) are EXD2, MAVS, OCIAD1 and TDRKH, all OMM proteins. The 32 preys only found in BioID (Mito-ER) are indicated. The 5 preys in bold were previously reported to have a function at mitochondria-ER contact sites (Csordas et al., 2018).

**Supplementary Figure S6** (related to Fig. 6)

**A.** Identification of several isoforms/processing products of EXD2 in human fibroblasts. Control human fibroblasts, untreated or treated with siEXD2 were analyzed by immunoblot and probed for the endogenous protein using EXD2 antibody. *α*-tubulin was used as a loading control.

**B.** Mitochondrial localization of EXD2-BirA* isoforms in Flp-In T-REx 293 cells. FLAG antibody was used to detect the EXD2 isoforms, TOMM20 was used as a mitochondrial marker. DAPI in blue. Scale bar, 10 μm.

**C.** EXD2 iso1 and M126A are membrane spanning proteins, while EXD2 iso2 is membrane attached/soluble protein. Alkaline carbonate extraction of mitochondria isolated from Flp-In T-REx 293 cells stably expressing EXD2-BirA*-FLAG-tagged isoforms was performed. Soluble proteins, or proteins attached to the membrane are found in the supernatant, while membrane-spanning proteins appear in the pellet. Antibodies against the indicated markers were used for comparison. FLAG antibody was used to visualize the EXD2 isoforms.

**D.** Pairwise comparison of EXD2-iso1 vs. EXD2-iso2 BioID results (log_2_ transformed average spectral counts normalized to total abundance). The majority of common preys (in blue) interact more strongly with EXD2-iso2, as seen by the number of spectra detected. Specific interactors in beige represent preys, where no spectra were detected for the other bait, while specific interactors in light grey passed the BFDR*≤*1% only for one bait.

**Supplementary Figure S7** (related to Fig. 7)

**A, B.** Enlarged views of mitochondrial ribosomal clusters detected in Fig. 7A. Several known, or predicted proteins involved directly or indirectly in mitochondrial ribosome assembly also cluster with mitochondrial ribosomal proteins (TRMT10C, FASTKD5, ERAL1, LRPPRC, MRM3, GUF1, MALSU1, GTPBP10, DDX28). (**A)** Two clusters of mtLSU and an mtSSU cluster. Proteins of the mtSSU are indicated in turquoise, proteins of the mtLSU-cluster 1 are shown in green, mtLSU-cluster 2 in dark blue and MRPL12 and MRPL53 in light green. (**B)** mtLSU-cluster 3; mtLSU proteins are shown in pink.

**Supplementary Table S1** (related to Fig. 1)

BioID proximity interactome for the matrix and IMS “environment”. SAINTexpress Tasks ID 3962 and 3961.

Associated files can be found in MassIVE submission MSV000085154 and PXD018196.

**Tab A.** Matrix MTS-BirA* dataset.

Bait indicates the MTS-“gene name” (column A); Prey (column B) is the NCBI protein accession number, PreyGene (column C) is the Symbol as per NCBI Gene; Spectral counts for each prey (column D, separated by “I” delimiter; column E, the sum of spectral counts; column F, the average of spectral counts), number of replicates performed (column G), spectral counts for the prey across all negative controls (column H), Averaged probability across replicates (AvgP; column I), maximal probability (column J), SAINT score (probability of protein-protein interaction, column K), Fold Change (counts in the purification divided by counts in the controls plus a small factor to prevent division by 0; column L) and Bayesian FDR (column M), Prey

Sequence Length (column N) are listed for each bait-prey relationship from the SAINTexpress output.

Note that the following entries were also manually removed from the final high confidence list associated with this manuscript: 1) “DECOY” sequences (removed during SAINTexpress); 2) all non-human protein contaminants (listed in **Tab C**).

**Tab B.** IMS MTS-BirA* dataset. Individual columns are identical as in **Tab A**.

Note that the following entries were also manually removed from the final high confidence list associated with this manuscript: 1) “DECOY” sequences (removed during SAINTexpress); 2) all non-human protein contaminants (listed in **Tab C**); 3) bait-bait interactions missed by SAINT: AIFM1 from MTS-AIFM1.

**Tab C**. List of removed non-human contaminants. Individual columns are identical as in **Tab A**. **Tab D.** List of removed entries that are no longer in NCBI. Individual columns are identical as in **Tab A**.

**Supplementary Table S2** (related to Fig. 1C-D)

Bait-prey enrichment over the matrix or IMS “environments”.

**Tab A.** Matrix-BirA* score for detected preys (matrix “environment”).

267 preys detected as significant proximity interactors for at least one of the three matrix MTS-BirA* baits (column A): dark brown - preys detected as significant for all three matrix MTS-BirA* baits; medium brown - preys detected as significant for two matrix MTS-BirA* baits; light brown - preys detected as significant for only one matrix MTS-BirA* bait.

MTS-OTC (column B): Prey-length normalized spectral counts for each prey normalized to total spectra detected for MTS-OTC-BirA*; MTS-COX8 (column C): Prey-length normalized spectral counts for each prey normalized to total spectra detected for MTS-COX8-BirA*; MTS-COX4 (column D): Prey-length normalized spectral counts for each prey normalized to total spectra detected for MTS-COX4-BirA*; Matrix-BirA* Score (column E): “average” of the calculated spectral counts in columns B-D (see methods for “average” calculation).

**Tab B.** IMS-BirA* score for detected preys (IMS “environment”).

120 preys detected as significant proximity interactors for at least one of the two IMS MTS-BirA* baits (column A): red-preys detected as significant for both IMS MTS-BirA* baits; orange - preys detected as significant for only one IMS MTS-BirA* bait.

MTS-AIFM1 (column B): Prey-length normalized spectral counts for each prey normalized to total spectra detected for MTS-AIFM1-BirA*; MTS-OPA1 (column C): Prey-length normalized spectral counts for each prey normalized to total spectra detected for MTS-OPA1-BirA*; IMS-BirA* Score (column D): “average” of the calculated spectral counts in columns B-C (see methods for “average” calculation).

**Tab C.** Enrichment score for all preys detected as significant proximity interactors of LRPPRC (as bait). Bait indicates the bait gene name (column A); Prey (column B) is the NCBI protein accession number; PreyGene (column C) is the Symbol as per NCBI Gene; NormSpec (column D), prey-length normalized spectral counts for each prey normalized to total spectra detected for LRPPRC as bait; Fold (column E), ratio between NormSpec (column D) and the normalized score for Matrix-BirA* (TabA, column E) for each prey; log2(fold), log2 of Fold (column E); N/A - prey not detected as a significant prey for any Matrix-BirA* bait.

**Tab D**. Enrichment score for all preys detected as significant proximity interactors of ACAD9 (as bait). Individual columns are identical as in **Tab C**.

**Tab E**. Enrichment score for all preys detected as significant proximity interactors of MTFR1L (as bait). Individual columns are identical as in **Tab C**.

**Tab F**. Enrichment score for all preys detected as significant proximity interactors of DIABLO (as bait). Bait indicates the bait gene name (column A); Prey (column B) is the NCBI protein accession number; PreyGene (column C) is the Symbol as per NCBI Gene; NormSpec (column D), prey-length normalized spectral counts for each prey normalized to total spectra detected for DIABLO as bait; Fold (column E), ratio between NormSpec (column D) and the normalized score for IMS-BirA* (TabA, column E) for each prey; log2(fold), log2 of Fold (column E); N/A - prey not detected as a significant prey for any IMS-BirA* bait.

**Tab G**. Enrichment score for all preys detected as significant proximity interactors of SLC25A12 (as bait). Individual columns are identical as in **Tab F**.

**Supplementary Table S3** (related to Fig. 2) Bait description table.

Bait Name contains a gene name, and if necessary, isoform and tag-localization (N- or C-terminal, column A). Bait Gene is the NCBI Entrez Gene (column B). Column C (BirA tag (N/C)) shows the tag localization for each bait, N-terminus/C-terminus. Uniprot ID, column D; Uniprot Entry Name, column E; Protein names, column F; Length, number of amino acids (column G), Mass, in Dalton (column H); Subcellular localization is based on Uniprot annotation (column I), Gene names, alternative gene names separated by space (column J). The presence of the bait in other mitochondrial databases (Mitocarta 2.0, IMPI, Gene Ontology) is shown in columns K-M.

**Supplementary Table S4** (related to Figs. 2-7)

High-confidence BioID proximity interactome for the mitochondrial network. SAINTexpress Task ID 3963.

Associated files can be found in MassIVE submission MSV000085154 and PXD018196.

**Tab A.** Bait indicates the bait gene name, isoform and the orientation of BirA*-FLAG tag (N or C; column A); Prey (column B) is the NCBI protein accession number, PreyGene (column C) is the Symbol as per NCBI Gene; Spectral counts for each prey (column D, separated by “I” delimiter; column E, the sum of spectral counts; column F, the average of spectral counts), number of replicates performed (column G), spectral counts for the prey across all negative controls (column H), Averaged probability across replicates (AvgP; column I), maximal probability (column J), SAINT score (probability of protein-protein interaction, column K), Fold Change (counts in the purification divided by counts in the controls plus a small factor to prevent division by 0; column L) and Bayesian FDR (column M), Prey Sequence Length (column N) are listed for each bait-prey relationship from the SAINTexpress output.

Note that the following entries were also manually removed from the final high confidence list associated with this manuscript: 1) “DECOY” sequences (removed during SAINTexpress); 2) all non-human protein contaminants (listed in **Tab B**); 3) mass-spec contaminants from previous samples and bait-bait interactions missed by SAINT (information listed in **Tab C**).

**Tab B.** List of removed non-human contaminants. Individual columns are identical as in **Tab A**. **Tab C**. List of preys either removed, or with an adjusted score due to a contamination from a previous mass-spec run. Two bait-bait interactions missed by SAINT are also listed. Sample number, sample number for the indicated mass-spec run in Prohits (column A); Bait, the bait gene name, isoform and the orientation of BirA*-FLAG tag (N or C; column B); Prey contaminant, Symbol as per NCBI Gene (column C), Action performed is indicated in column D.

**Tab D.** List of removed entries that are no longer in NCBI. Individual columns are identical as in

Tab A.

**Supplementary Table S5** (related to Fig. 2)

List of unique interactions between mitochondrial baits and their identified preys.

The preys identified in this list have been found interacting with a single mitochondrial bait with a BFDR*≤*1%. Bait indicates the bait gene name, isoform and the orientation of BirA*-FLAG tag (N or C; column A); Prey (column B) is the NCBI protein accession number, PreyGene (column C) is the Symbol as per NCBI Gene; Spectral counts for each prey (column D, separated by “I” delimiter; column E, the sum of spectral counts; column F, the average of spectral counts), number of replicates performed (column G), spectral counts for the prey across all negative controls (column H), Averaged probability across replicates (AvgP; column I), maximal probability (column J), SAINT score (probability of protein-protein interaction, column K), Fold Change (counts in the purification divided by counts in the controls plus a small factor to prevent division by 0; column L) and Bayesian FDR (column M), Prey Sequence Length represents the number of amino acids (column N), Average Ctrl Counts (column O) is the average spectral counts detected by negative controls (average of column H), Prey-length Normalized Spectral Counts (column P). Column Q shows if the prey-bait interaction exhibits infinite specificity, signifying that no spectra were detected for the prey by any other bait (Y-yes/N-no).

**Supplementary Table S6** (related to Fig. 5)

**Tab A.** List of identified GO terms: Cellular Compartment for all non-Mitocarta preys detected by OMM baits.

Bait indicates the bait gene name, isoform and the orientation of BirA*-FLAG tag (N or C; column A); GO term Cellular Compartment based on PANTHER (GO ontology database Released 2019-02-02, column B), Fold enrichment score was assigned to each GO term based on the fold enrichment score in PANTHER (column C). The most specific subclass terms are indicated. All GO:terms were identified with BFDR*≤*1%.

**Tab B.** List of proteins classified as mitochondia-ER contact site proteins used for generation of **Supplementary Figure S5D**. BioID (Mito-ER) proteins represent 72 preys common to the cluster of five OMM baits (SLC25A46, PTPN1, FKBP8, RMDN3 and RHOT2). The lists of previously published proteomic datasets of mitochondria-ER contact site proteins were used for comparison: (Hung et al., 2017)(Cho et al. (https://doi.org/10.1101/2020.03.11.988022) (Kwak, 2020).

**Supplementary Table S7** (related to Fig. 6)

BioID proximity interactome of EXD2 isoforms. SAINTexpress Task ID 3963. Associated files can be found in MassIVE submission MSV000085154 and PXD018196.

**Tab A.** Bait indicates the bait gene name and an isoform (column A); Prey (column B) is the NCBI protein accession number, PreyGene (column C) is the Symbol as per NCBI Gene; Spectral counts for each prey (column D, separated by “I” delimiter; column E, the sum of spectral counts; column F, the average of spectral counts), number of replicates performed (column G), spectral counts for the prey across all negative controls (column H), Averaged probability across replicates (AvgP; column I), maximal probability (column J), SAINT score (probability of protein-protein interaction, column K), Fold Change (counts in the purification divided by counts in the controls plus a small factor to prevent division by 0; column L) and Bayesian FDR (column M), Prey Sequence Length (column N) are listed for each bait-prey relationship from the SAINTexpress output.

Note that the following entries were also manually removed from the final high confidence list associated with this manuscript: 1) “DECOY” sequences (removed during SAINTexpress); 2) all non-human protein contaminants (listed in **Tab B**); 3) mass-spec contaminant (CLUH) from a previous sample run for EXD2_iso1 sample run #9416 was also removed.

**Tab B.** List of removed non-human contaminants. Individual columns are identical as in **Tab A**. **Tab C.** List of removed entries that are no longer in NCBI. Individual columns are identical as in **Tab A**.

**Supplementary Table S8**

List of clones and primers used for bait-BirA* cloning.

Bait Name contains a gene name, and if necessary, isoform and tag-localization (N- or C-terminal, column A). Column B (BirA* tag (N/C)) shows the tag localization for each bait, N-terminus/C-terminus. Uniprot ID, column C; clone#, column D; Source supplying the clone, column E. Toronto - signifies construct was obtained from the Lunenfeld-Tanenbaum Research

Institute (LTRI) Open Freezer; PCR primers (columns F and G), PCR primers used for amplification of individual genes; Cloning strategy, column H.

## METHODS

### LEAD CONTACT AND MATERIALS AVAILABILITY

Further information and requests for resources and reagents should be directed to and will be fulfilled by the Lead Contact, Eric A. Shoubridge.

### EXPERIMENTAL MODEL AND SUBJECT DETAILS

#### Mammalian cell culture

Flp-In™ T-REx™ 293 cells (Invitrogen), Phoenix packaging cell line (a kind gift of Garry P. Nolan) and control human fibroblasts (Montreal Children Hospital Cell bank) were grown in high-glucose DMEM (Wisent) supplemented with 10% fetal bovine serum, 500 units/ml penicillin, and 500 µg/ml streptomycin in 5% CO_2_ incubator at 37°C. Cell lines were regularly tested for mycoplasma contamination. For immunofluorescence analysis, cells were cultured on 12 mm round coverslips (thickness No.1, Fisherbrand) in 24-well plates.

### Flp-In™ T-REx™ 293 stable cell line generation

Flp-In™ T-REx™ 293 cells were seeded at 250,000 cells per well in a 6-well plate in 2 ml of media. The next day, cells were transfected using Lipofectamine 2000 (Invitrogen) with 200 ng pDEST-pcDNA5-ProteinX-BirA*-FLAG, and 2 μg of pOG44 in 250 μl of 1x Opti-MEM (Invitrogen) mixed with 5 μl of Lipofectamine 2000 reagent in 250 μl of 1x Opti-MEM. The Opti-MEM/Lipofectamine solution was added to the Opti-MEM/plasmids solution and incubated 20 minutes before addition to the cells (in media without antibiotics). The media was changed 4 hours after transfection. The next day, transfected cells were passaged to 10 cm plates and the following day selected by the addition of hygromycin (Wisent) to the growth media at a final concentration of 200 μg/ml. This selection media was changed every 2-3 days until clear visible colonies were present. Up to 6 colonies per construct were picked, expanded until further analysis.

### mCherry-ER overexpression cell line generation

Control human fibroblast line stably over-expressing mCherry-ER marker was created by retroviral overexpression of pBabe-mCherry-ER. pBabe-mCherry-ER plasmid was transiently transfected into Phoenix packaging cell line using HBS/Ca_3_(PO_4_)_2_ method (https://web.stanford.edu/group/nolan/_OldWebsite/publications/publications.html). Phoenix cells were seeded on a 6 cm plate at about 40-50% confluency one day before transfection. On the day of transfection, 3 ml of fresh media were added to the cells and 25 μM chloroquine was added to the media 5 minutes prior to transfection. pBabe-mCherry-ER plasmid (2 μg) supplemented with 3 μg ssDNA was diluted to a final volume of 0.439 ml and 61 μl of 2M CaCl_2_ was added to the DNA mixture. Under constant, vigorous bubbling, 0.5 ml of 2x HEPES-buffered saline (2xHBS, pH 7.0) was added to the DNA/CaCl_2_ solution. HSB/DNA mixture was then added dropwise to the Phoenix cells. Day after, Phoenix cells were detoxified by media replacement, and the produced virus was collected for the following 24 hours. Virus-containing media collected from Phoenix cells were filtered through a 0.45 μM syringe filter, polybrene was added to a final concentration of 4 μg/ml and 1 ml was added to a 6 cm plate of 70% confluent control human fibroblasts. The next day, fibroblasts were passaged to a 10 cm plate and selected by addition of 2 µg/ml of puromycin.

### METHOD DETAILS

#### Bait selection and cloning

Mitochondrial baits were chosen to represent all sub-mitochondrial compartments and we chose both confirmed and predicted mitochondrial proteins. All matrix, IMS and most IMM baits were tagged at the C-terminus, as the N-terminus is essential for their mitochondrial targeting. One IMM protein (SLC25A12) was tagged at the N-terminus, and one IMM protein (SLC25A51) was tagged at both termini. For OMM proteins, the position of the tag was either inferred from literature; was based on the protein structure profiling using a transmembrane prediction program TMHMM (http://www.cbs.dtu.dk/services/TMHMM/); or tagging of both ends was tested. Finally, nine OMM proteins were tagged at the N-terminus, four at the C-terminus and six proteins were tagged at both N- and C-terminus. Overall, 93 different proteins were profiled, including 4 proteins, where two different isoforms were represented. All constructs and the position of their BirA*-FLAG tag are listed in the **Supplementary Table S3**.

All constructs were generated using Gateway cloning into an appropriate pDEST-pcDNA5-BirA*-FLAG construct (to create in-frame N- or C-terminal BirA*-FLAG fusion proteins). Selected entry Gateway clones were obtained from DNASU, Addgene or the Lunenfeld-Tanenbaum Research Institute (LTRI) Open Freezer (Toronto, Canada). If no entry clone was commercially available, entry clones were created by PCR amplification of the ORF of interest from human cDNA, followed by a sub-cloning into pDONR™-221 (Invitrogen). The EXD2-M126A variant was created by site-directed mutagenesis of pDEST-EXD2_iso1-BirA*-FLAG construct using Quick Change Lightning kit (Stratagene).

To interrogate the protein environment of the mitochondrial matrix and IMS we created MTS-targeted BirA* constructs, in which the MTS is cleaved off after the import into the appropriate space. MTS-COX8 (N-terminal 29 amino acids), MTS-AIFM1 (N-terminal 120 amino acids) and MTS-OPA1 (N-terminal 208 amino acids) were amplified by PCR from human cDNA and cloned into pDEST-pcDNA5-BirA*-FLAG-C-ter using Gateway cloning. For MTS-COX4 (N-terminal 22 amino acids) and MTS-OTC (N-terminal 32 amino acids), PCR primers encompassing the whole sequence plus BsrGI overhangs were annealed together and subsequently ligated into a BsrGI digested pDEST-pcDNA5-BirA*-FLAG-C-ter. The clones and primers used for cloning of all the individual constructs are listed in **Supplemental Table S8**. All final plasmids were verified by Sanger sequencing.

#### pBabe-mCherry-ER Cloning

mCherry-ER fusion protein was amplified by PCR from mCherry-ER-3 plasmid (Addgene) and cloned into Gateway modified pBABE (in house). The sequence was verified by Sanger sequencing.

#### Immunofluorescence

For selection of stable Flp-In™ T-REx™ 293 expressing clones, the localization and the expression level of the construct was assessed by immunofluorescence using anti-FLAG antibody (Sigma). Expression of the construct was induced or not by addition of 1 μg/ml tetracycline (Sigma) for 24 hours in cells grown on coverslips. On the following day, cells were fixed using 4% formaldehyde in PBS for 20 min at 37°C. Coverslips were washed 3 times using PBS and cells were solubilized in 0.05% Triton in PBS for 15 min at room temperature. After three PBS washes, coverslips were blocked in PBS containing 5% BSA for 10 minutes, incubated with primary antibodies (anti FLAG 1:4000 and anti-TOMM20 1:2000) for one hour at room temperature, washed 3 times with PBS and incubated with Alexa conjugated secondary antibodies (1:2000) and DAPI (1:2000) for 30 min at room temperature. Coverslips were washed in PBS (3 times), mounted and cells were imaged with Olympus IX83 microscope connected with Yokogawa CSU-X confocal scanning unit, using UPLANSAPO 100x/1.40 Oil objective (Olympus) and Andor Neo sCMOS camera. Images were processed in Fiji (Schindelin et al., 2012). For clone selection, predominant mitochondrial localization was required, as well as minimal expression of the construct in non-induced cells. One clone per construct was selected and used for BioID analysis. The representative immunofluorescence images for each bait are available on ProHits-web.lunenfeld.ca.

For confirmation of the localization of selected proteins, control human fibroblasts were used. The immunocytochemistry was performed as mentioned above, with the exception of cell solubilization, where 0.1% Triton was used. Anti-PDH (1:1000) or anti-cytochrome c (1:1000) antibodies were used to visualize mitochondria, ABCD3 (1:1000) was used as a peroxisomal marker and ER was visualized using cells stably overexpressing mCherry-ER (ER-red). Appropriate Alexa conjugated secondary antibodies (1:2000) were used and cells were imaged as mentioned above.

#### BioID sample preparation

The selected Flp-In™ T-REx™ 293 clones and BioID control cells (Flp-In™ T-REx™ 293 cells alone or Flp-In™ T-REx™ 293 over-expressing BirA*-FLAG-GFP) were scaled up to the needed number of 150 mm plates - one to freeze back and 6 for treatments and harvesting. Cells were grown to 70% confluency before induction of protein expression using 1 μg/ml tetracycline (Sigma), and media supplementation with 50 μM biotin for protein labeling. Cells were harvested 24 hours later as follows: cell media was decanted; cells were washed twice with 5 ml PBS per 150 mm plate and then harvested by scraping in 5 ml of PBS. Cells from 3 x 150 mm plates were pelleted at 800 rpm for 3 minutes, PBS aspirated, and pellets transferred to −80°C freezer. A small fraction (2%) of each sample, including BioID control cells, was set aside to be tested for induction of the bait expression and biotinylation by SDS-PAGE/Western-blotting using anti-FLAG and anti-biotin antibodies.

Cell pellets (corresponding to 3 x 150mm plates) were incubated at 4°C in RIPA buffer (50 mM Tris-HCl pH 7.5, 150 mM NaCl, 1% NP-40, 1mM EDTA, 1 mM EGTA, 0.1% SDS, Sigma protease inhibitors 1:500, and 0.5% Sodium deoxycholate) at 1:10 (pellet weight in g: lysis buffer volume in ml) for 1 hour on a nutator. After incubation, 1 μl of benzonase (250U) was added to each sample and the lysates sonicated (3 x 10 second bursts with 2 seconds rest in between) on ice at 65% amplitude. These lysates were then centrifuged for 30 min at 20,817g at 4°C. After centrifugation supernatants were transferred to new 15 ml falcon tubes and a 30 μl bed volume of pre-washed streptavidin-sepharose beads (GE) were added to each sample. The streptavidin-sepharose beads were pre-washed 3 times with 1 ml RIPA buffer (minus protease inhibitors and sodium deoxycholate); beads were pelleted at 400g, 1 minute in-between washes. Affinity purification was performed at 4°C on a nutator for 3 hours. After the purification, the beads were pelleted (400g, 1 min), the supernatant removed, and the beads transferred to a 1.5 ml microcentrifuge tube in 1 ml RIPA buffer (minus protease inhibitors and sodium deoxycholate). The beads were then washed by pipetting up and down (4x per wash step) first with an additional 1 ml RIPA buffer (minus protease inhibitors and sodium deoxycholate) followed by two washes in TAP lysis buffer (50 mM HEPES-KOH pH 8.0, 100 mM KCl, 10% glycerol, 2 mM EDTA, 0.1% NP-40) and then three washes in 50 mM ammonium bicarbonate pH 8. Beads were pelleted by centrifugation (400g, 1 min) and the supernatant aspirated in between wash steps. After the last wash all residual 50 mM ammonium bicarbonate was pipetted off and proteins digested on bead.

After affinity purification and removal of all washing buffer, streptavidin-sepharose beads were resuspended in 200 µl of 50 mM ammonium bicarbonate pH 8 containing 1 µg of trypsin (Sigma-Aldrich). Samples were then incubated at 37°C overnight with mixing on a rotating disc. The next day an additional 0.5 µg of trypsin was added to each sample (in 10 µl 50 mM ammonium bicarbonate pH 8) and samples incubated for an additional 2 hours at 37 °C with mixing on a rotating disk. Beads were then pelleted at 400g for 2 min and the supernatant transferred to a fresh 1.5 ml microcentrifuge tube. The beads were rinsed 2 times with 150 µl of 50 mM ammonium bicarbonate pH 8 (pelleting beads at 400g, 2 min in between) and these rinses were combined with the original supernatant. The pooled supernatant was then centrifuged at 16,100g for 10 min and most of the supernatant (minus 30 µl to get rid of all beads) was transferred to a new 1.5 ml microfuge tube. These samples were then dried in a centrifugal evaporator.

#### Mass-Spec data acquisition

Affinity-purified, digested material from 3 x 150 mm plates was resuspended in 12 μl of 5% formic acid and centrifuged at 16,100g for 1 min before 5 μl was injected by autosampler to a home-made HPLC column (0.75 μm ID, 350 μm OD with spray tip generated using a laser puller) loaded with 10 to 12 cm of C18 reversed-phase material (ZorbaxSB, 3.5 μm or ReproSil-Pur 120 C18-AQ 3μm). The column was placed in-line with a LTQ-Orbitrap Velos (Thermo Fisher Scientific) equipped with a nanoelectrospray ion source (Proxeon, Thermo Fisher Scientific) connected in-line to a NanoLC-Ultra 2D plus HPLC system (Eksigent, Dublin, USA). The LTQ-Orbitrap Velos or Elite instrument under Xcalibur 2.0 was operated in the data dependent mode to automatically switch between MS and up to 10 subsequent MS/MS acquisition. Buffer A is 99.9% H_2_O, 0.1% formic acid; buffer B is 99.9% acetonitrile, 0.1% formic acid. The HPLC gradient program delivered an acetonitrile gradient over 125 min. For the first 20 min, the flow rate was of 400 μl/min at 2% B. The flow rate was then reduced to 200 μl/min and the fraction of solvent B increased in a linear fashion to 35% until 95.5 min. Solvent B was then increased to 80% over 5 min and maintained at that level until 107 min. The mobile phase was then reduced to 2% B until the end of the run (125 min).

#### Proteinase K assay

Flp-In™ T-REx™ 293 cells stably expressing EXD2-iso1-BirA*-FLAG, EXD2-iso2-BirA*-FLAG or EXD2-M126A-BirA*-FLAG were grown on 3×15cm plates and the expression of the constructs was induced for 24 hours by addition of 1 μg/ml tetracycline. Cells were washed twice with PBS, scraped in ice-cold PBS and pelleted at 800 rpm for 3 minutes. Cell pellets were resuspended in 800 µl of MIB buffer (220 mM mannitol, 68 mM sucrose, 10 mM HEPES pH 7.4, 80 mM KCl, 0.5 mM EGTA, 2mM magnesium acetate) and homogenized by ∼15 strokes of teflon-glass homogenizer at 2100rpm. Homogenized cells were spun twice at 600g for 5 minutes to remove nuclei and unbroken cells. Mitochondria were pelleted by centrifugation at 10000g for 10 min, and washed twice with MIB buffer. 20 µg of mitochondria (at a final concentration 1 mg/ml in MIB buffer) were either left untreated, or were treated for 30 min on ice with 50 µg/ml of proteinase K alone or in combination with 1% Triton-X100. After the proteinase K treatment, 2 µl of 40 mM phenylmethanesulfonyl fluoride (PMSF) was added to all samples and samples were incubated on ice for 20 min. Following the protease inhibition, Laemmli loading buffer was added and samples were boiled for 5 min before separation by SDS-PAGE.

#### Alkaline carbonate extraction

Mitochondria were prepared as mentioned above for proteinase K assay from Flp-In™ T-REx™ 293 cells stably expressing EXD2-iso1-BirA*-FLAG, EXD2-iso2-BirA*-FLAG or EXD2-M126A-BirA*-FLAG. 50 µg of mitochondria were resuspended in 100 µl of 100 mM sodium carbonate buffer (pH 11.5) and incubated on ice for 30 min with a gentle mixing by pipetting every 10 min. 50 µl of alkaline carbonate treated mitochondria were set aside as input (kept on ice) and 50 µl were centrifuged at 100 000g for 30 min at 4°C in TLA100 rotor (Beckman Coulter). Pellet was gently washed with ice-cold sodium carbonate buffer and resuspended in 20 µl of 1x Laemmli loading buffer. Supernatant and input were further precipitated by TCA (trichloroacetic acid). 8.8 µl of 100% (w/v) TCA was added to 50 µl of sample and sample was stored at −80°C overnight. TCA precipitated proteins were centrifuged at 20 000g for 20 min at 4°C, washed twice with 500 µl of ice-cold acetone, and air-dried. Final TCA precipitates were resuspended in 20 µl of 1x Laemmli loading buffer.

#### SDS-PAGE and Western blot

Proteinase K assay samples, alkaline carbonate samples, control and siRNA treated cells were separated by 10% Laemmli SDS-PAGE electrophoresis. Gels were transferred to a nitrocellulose membrane (PALL), and subsequently incubated with indicated primary and secondary antibodies in 5% skim-milk Tris-buffered saline solution with 0.1% Tween 20.

#### siRNA treatment

Control human fibroblasts were transfected with 12 nM EXD2 siRNA (Dharmacon) using 2.5 µl Lipofectamine RNAiMAX (Invitrogen) according the manufacturer’s instructions using a reverse transfection procedure in a 6-well plate. The treatment was repeated on day 3 and cells were harvested for SDS-PAGE on day 6.

### QUANTIFICATION AND STATISTICAL ANALYSIS

#### MS data analysis for BioID

All Thermo RAW files were saved in our local interaction proteomics LIMS, ProHits (Liu et al., 2016). mzXML files were generated from Thermo RAW files using the ProteoWizard (Adusumilli and Mallick, 2017) converter, implemented within ProHits (--filter “peakPicking true2” --filter “msLevel2”). The searched database contained the human and adenovirus complement of the RefSeq protein database (version 57) supplemented with “common contaminants” from the Max Planck Institute (http://141.61.102.106:8080/share.cgi?ssid=0f2gfuB) and the Global Proteome Machine (GPM; http://www.thegpm.org/crap/index.html) and common sequence tags. The sequence database consisted of forward and reversed sequences; in total 72,226 sequences were searched. The search engines were Mascot and Comet, with trypsin specificity and two missed cleavage sites allowed. Methionine oxidation and asparagine/glutamine deamidation were set as variable modifications. The fragment mass tolerance was 0.6 Da and the mass window for the precursor was ±12 ppm. The resulting Comet and Mascot search results were individually processed by PeptideProphet (Keller et al., 2002) and peptides were assembled into proteins using parsimony rules first described in ProteinProphet (Nesvizhskii et al., 2003) into a final iProphet (Shteynberg et al., 2011) protein output using the Trans-Proteomic Pipeline (TPP; Linux version, v0.0 Development trunk rev 0, Build 201303061711). TPP options were as follows. For the Velos Orbitrap files, general options are -p0.05 -x20 -PPM - d“DECOY”, iProphet options are pPRIME and PeptideProphet options are pPAEd. All proteins with a minimal iProphet protein probability of 0.05 were parsed to the relational module of ProHits. Note that for analysis with SAINT, only proteins with iProphet protein probability ≥ 0.95 were considered. This corresponded to an estimated protein level FDR of ∼0.5%. A minimum of two detected peptide ions was also enforced.

#### SAINT analysis

SAINT (Significance Analysis of INTeractome) calculates, for each prey identified in an experiment, the probability of true interaction by using spectral counts (semi-supervised clustering, using a number of negative control runs). SAINTexpress (Teo et al., 2014) analysis was performed using version exp3.6.1. Three SAINTexpress analyses were performed: (1) SAINT #3963: contained all mitochondrial baits as well as EXD2-iso2 and EXD2-M126A and the analysis was performed with two biological replicates per bait; (2) SAINT #3962: contained MTS-OPA1 and the analysis was performed with two biological replicates per bait; (3) SAINT #3961: contained MTS-COX8, MTS-COX4, MTS-OTC, MTS-AIFM1 and the analysis was performed with 6-8 biological replicates per bait compressed to 6 replicates. Bait protein samples for all three SAINT analyses were analyzed alongside 48 negative control runs, consisting of 24 purifications from untransfected Flp-In™ T-REx™ 293 cells and 24 purifications from cells expressing BirA*-FLAG-GFP, compressed to 24 as previously described, to increase the stringency of the scoring (Mellacheruvu et al., 2013). For downstream analysis, the three SAINT files were combined/separated to create 4 datasets: **Supplementary Table S4:** contains the core 100 mitochondrial baits, **Supplementary Table S1:** contains MTS-BirA* data in two sets: Matrix-BirA* (data from MTS-COX8, MTS-COX4 and MTS-OTC) and IMS-BirA* (data from MTS-OPA1 and MTS-AIFM1) and **Supplementary Table S7:** contains data on EXD2-iso1 (same data as in **Supplementary Table S4** for this bait), EXD2-iso2 and EXD2-M126A. All non-human protein contaminants were removed from the SAINT files. DHX30-DHX30 and METTL17-METTL17 bait-prey interactions were also manually removed. Entries no longer in NCBI were also manually removed. Three entries were updated based on new NCBI annotation, namely LOC100507855 to AK4, LOC101060541 to RBM8A and LOC101060751 to MRC1. **Supplementary Tables S1, S4 and S7** show the entire SAINT analysis and a list of preys either removed, or with an adjusted score as a result of an aberrant score due to a contamination from a previous mass-spec run. A 1% Bayesian false discovery rate (BFDR) cutoff was used to select confident proximity interactors for downstream analysis. The SAINT #3963 dataset is also readily available on ProHits-web.lunenfeld.ca for interactive exploration.

#### Databases used for analysis

Mitocarta 2.0 (Calvo et al., 2016) was used for annotation of detected preys as mitochondrial or non-mitochondrial, with the consideration that 20 different proteins have updated gene names since the publication of Mitocarta 2.0 database (Mitocarta 2.0 names in parenthesis): RCC1L (WBSCR16), SDHAF3 (ACN9), MRM3 (RNMTL1), MRPL58 (ICT1), TWNK (C10orf2), GATB (PET112), TOMM70 (TOMM70A) and 13 subunits of mitochondrial ATP synthase. Human gene annotations were downloaded from Gene Ontology (GO) on 17/04/2019 (GO version date 02/02/2019). The Human Protein Atlas (HPA) subcellular localization data was downloaded on 06/06/2019 from www.proteinatlas.org/about/download and is based on the Human Protein Atlas version 18.1.

#### Data analysis and visualization using ProHits-viz

The SAINT files presented in **Supplementary Tables S1, S4 and S7** were used as input files for ProHits-viz (Knight et al., 2017) analysis using the modules and parameters indicated below.

**Bait-bait heatmap**: the SAINT output file from **Supplementary Table S4** was uploaded to ProHits-viz Dot plot generator module, which was used with default settings, except that the distance metric was set to Jaccard. The Jaccard distance is defined as 1 – J(A,B), where J(A,B) is the Jaccard index, defined as the overlap between two sets (A, B) and is calculated as 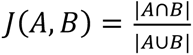. The dendrogram was automatically created in the module. In the final figure, labelling of the mitochondrial compartments was performed manually outside of ProHits-viz.

**Bait vs Bait comparison**: each of the SAINT output files was uploaded to ProHits-viz Bait-Bait Comparison module individually. The reference (x axis) and secondary (y axis) baits were selected and the “versus” output type was selected. Both the primary and the secondary filter values were set at BFDR*≤*1% (0.01). Subtraction of the spectral counts across the controls and normalization to total abundance value were performed and the data were log2 transformed. On these figures, the edge circle color is mapped to the BFDR for the indicated prey across both baits compared. A full black circle indicates that the BFDR*≤*1% threshold has been met with both baits, while a half circle indicates that this cutoff has been reached with only one of the baits. Note, that SAINT removes bait-bait interactions; thus, we color-coded the baits in bait-bait comparisons when detected as a prey. Color-coding and labeling of selected preys were performed manually outside of ProHits-viz.

**Specificity plots**: the SAINT output file from **Supplementary Table S4** was uploaded to ProHits-viz Prey Specificity module and Fold Enrichment analysis was performed. Subtraction of the spectral counts across the controls was performed. The spectral counts for each prey were normalized to the Prey Sequence Length. The figures were manually annotated and color-coded outside of ProHits-viz.

**Correlation analysis**: the SAINT output file from **Supplementary Table S4** was uploaded to the ProHits-viz Correlation Analysis module. The abundance measure was set to AvgSpectra for **Fig. 4, 5 and S4** and to FoldChange for Fig. 7 **and S7**. The score filter value was set at BFDR*≤*1% (0.01). Abundance cutoff of 5 spectra, minimum bait requirement of 2 and subtraction of the counts in the controls were performed. Pearson correlation with the Euclidian metric was used, with “complete” clustering. Calculation of GO term enrichment was performed directly in ProHits-viz, through the isolation of boxed areas using the dynamic viewer, and calculation of term enrichment through g:Profiler (Reimand et al., 2016) using all default options). Annotation of the GO terms was subsequently done manually outside of ProHits-viz. Some panels were zoomed in from the dynamic viewer.

**Heat Map generation**: **Supplementary Table S6** was uploaded to ProHits-viz Dot plot generator module, which was used with default settings, with a secondary filter set to 0.01. The resulting dot plot was visualized as a heat map in the dynamic viewer “display options”. The dendrogram was automatically created in the module. For the **Supplementary Table S6**, all non-mitochondrial preys detected by OMM baits were uploaded to PANTHER and GO term Cellular Compartment analysis was performed. All GO terms were identified with BFDR*≤*1%. The most specific subclass terms and their Fold enrichment score were used for further analysis. The resulting figure was color-coded and labelled manually outside of ProHits-viz.

#### Data analysis and visualization using Cytoscape

Cytoscape 3.7.2 (Shannon et al., 2003) was used. For the generation of the network in Fig. 7, a correlation matrix data from the prey-prey analysis was used. Only correlations greater than 0.7 were considered for the network, and only edges with at least one mitochondrial ribosomal protein were included. The default layout was selected. The edges are color-coded based on the correlation score. Node size shows the degree of connectivity of each node, and is defined as a number of edges linked to a node. The layout was exported as a PDF, and manually adjusted: mtSSU proteins are indicated in turquoise and light blue; and the mtLSU proteins are in dark blue, green and pink. Other interactors, which are not part of mitochondrial ribosome, are indicated in grey.

For the generation of the bait network images on ProHits-web.lunenfeld.ca, the bait-prey proximity network restricted to the 25 top proximity interactors for each bait (see details below) was used. For all individual networks, all edges connected to the bait of interest were first selected alongside all the nodes connected to these edges. Network was then created from selected nodes, all edges. yFiles Organic Layout was selected with the length-normalized spectral counts (NormSpec) used as edge weight. NormSpec was used for continuous mapping of the edge width, and the directional network was created (displaying bait → prey relationships). Node size indicates the frequency of detection of the indicated protein in the “top 25” network. Proteins used as baits in the dataset are indicated by a purple color. The presence of the protein in Mitocarta 2.0 is shown as a black border around the node. Nodes were manually re-arranged from this layout to increase visibility and highlight specific proximity interactions. The layout was exported as a PDF, manually adjusted, and eventually converted to a .png file which was uploaded to ProHits-web.lunenfeld.ca.

#### “Top 25” interactors

The top 25 interacting preys for each bait were determined from their length-normalized spectral counts (NormSpectra). For bait x and prey y the NormSpectra are calculated by first subtracting the prey’s average spectral count found in controls from its abundance with bait x, then multiplying by the median length of all other preys across bait x and dividing by its length:

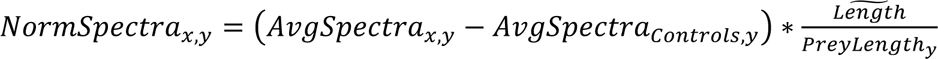

#### Specificity analysis of the mitochondrial “environment”

High-confidence preys detected by Matrix-BirA* and IMS-BirA* were compared to previously published datasets for mitochondrial matrix APEX (Rhee et al., 2013) and intermembrane space APEX (Hung et al., 2014). To calculate the mitochondrial specificity of the matrix and IMS proteome, we first determined if a prey had been annotated as a mitochondrial protein according to the Mitocarta 2.0 (Calvo et al., 2016) or Gene Ontology (GO) database. The number of preys with previous mitochondrial annotation was then divided by the total number of detected preys. Matrix specificity and “IMS exposure” were calculated as described in MTS-APEX and IMS-APEX: Matrix-BirA* and MTS-APEX were compared to a dataset with 424 proteins with sub-mitochondrial localization (Rhee et al., 2013). IMS-BirA* and IMS-APEX were compared to a dataset with 579 proteins with sub-mitochondrial localization (Hung et al., 2014). Sub-mitochondrial annotation was available for 68 IMS-BirA* detected preys and for 142 Matrix-BirA* detected preys, respectively. Proteins were considered “IMS exposed” if they were previously annotated as OMM, IMS or IMM proteins.

#### Calculation of the Matrix-BirA* score

The prey-length normalized spectral counts for each of the 267 high-confidence preys (BFDR*≤*1%) identified by three mitochondrial Matrix-BirA* baits were normalized to the total spectra detected by each Matrix-BirA* bait (Norm2Spec count). The Matrix-BirA* Score was then calculated as an average of the three Norm2Spec counts for preys identified by all three Matrix-BirA* baits, mean of two Norm2Spec counts for preys identified by two baits, and was made equal to the Norm2Spec count of the one bait, where prey was detected by one bait only.

#### Calculation of the IMS-BirA* score

The prey-length normalized spectral counts for each of the 120 high-confidence preys (BFDR*≤*1%) identified by two mitochondrial IMS-BirA* baits were normalized to the total spectra detected by each IMS-BirA* bait (Norm2Spec count). The IMS-BirA* Score was then calculated as a mean of the two Norm2Spec counts for preys identified by two IMS-BirA* baits, or was made equal to the Norm2Spec count of the one bait, where prey was detected by one bait only.

#### Visualization of proteins of the mitochondrial ribosome

The cryo-EM structure of the human mitochondrial ribosome (PDB 3J9M, (Amunts et al., 2015) was loaded into the Swiss-PDBViewer and either the large mitochondrial ribosomal subunit, or the small mitochondrial ribosomal subunit were viewed individually. The structure was rendered in Solid 3D. The individual RNA molecules were color-coded in grey. The proteins were shown as ribbons and color-coded based on the individual clusters, in which they were identified in the prey-prey correlation analysis.

#### Other analysis

All other statistical analysis and data plotting was performed using Microsoft Excel and Prism 8 (GraphPad). Venn diagrams were created using either Venny 2.1 (https://bioinfogp.cnb.csic.es/tools/venny/index.html) or area-proportional Venn diagram software Biovenn (Hulsen et al., 2008).

### DATA AND CODE AVAILABILITY

All presented data is available in the main text and supplementary materials.

Data were also deposited to ProHits (http://prohits-web.lunenfeld.ca), where information on individual baits and their detected preys can be found.

Dataset consisting of raw files and associated peak lists and results files have been deposited in ProteomeXchange through partner MassIVE (http://proteomics.ucsd.edu/ProteoSAFe/datasets.jsp) as a complete submission. Additional files include the sample description, the peptide/protein evidence and the complete SAINTexpress output for each dataset, as well as a “README” file that describes the dataset composition and the experimental procedures associated with the submission.

Dataset: Antonicka_et_al_MitoMap_P129_2020 MassIVE ID MSV000085154 and PXD018196

## KEY RESOURCES TABLE

### Antibodies

| REAGENT | SOURCE | IDENTIFIER |
| --- | --- | --- |
| anti TOMM20 | Santa Cruz | Cat# SC-11415, RRID:AB_2207533 |
| anti FLAG | Sigma | Cat# F3165, RRID:AB_25952<br>Cat# F1804, RRID:AB_262044 |
| anti FKBP8 | Sigma | Cat#HPA045177, RRID:AB_2679244 |
| anti PDH | Abcam | Cat# ab110333, RRID:AB_10862029 |
| anti OCIAD1 | Sigma | Cat# HPA044803, RRID:AB_10961287 |
| anti ABCD3 | Sigma | Cat# SAB4200181, RRID:AB_10639362 |
| anti cytochrome c | BD Biosciences | Cat# 556432, RRID:AB_396416 |
| anti SLC25A46 | Proteintech | Cat# 12277-1-AP, RRID:AB_2239384 |
| anti MFN1 | Cell Signaling Technology | Cat# 14739, RRID:AB_2744531 |
| anti RMDN3 | Proteintech | Cat# 20641-1-AP, RRID:AB_10695187 |
| anti YME1L1 | Proteintech | Cat# 11510-1-AP, RRID:AB_2217459 |
| anti SCO1 | Leary SC, HMG 2004 | PMID: 15229189 |
| anti LRPPRC | Sasarman F, Mol Biol Cell 2010 | PMID: 20200222 |
| anti GRSF1 | Sigma | Cat# HPA036985, RRID:AB_10672785 |
| anti EXD2 | Sigma | Cat# HPA005848, RRID:AB_1078768 |
| anti LONP1 | Proteintech | Cat# 15440-1-AP, RRID:AB_2137152 |
| anti SDHA | Abcam | Cat# ab14715, RRID:AB_301433 |
| anti MT-CO2 | a gift by Nancy Kennaway |  |
| Goat Anti-Mouse IgG (H+L) Highly Cross-adsorbed Antibody, Alexa Fluor 488 Conjugated | Molecular Probes | Cat# A-11029, RRID:AB_138404 |
| Goat anti-Rabbit IgG (H+L) Highly Cross-Adsorbed Secondary Antibody, Alexa Fluor 594 | Molecular Probes | Cat# A-11037, RRID:AB_2534095 |
| Goat Anti-Mouse IgG2a Antibody, Alexa Fluor 647 Conjugated | Molecular Probes | Cat# A-21241, RRID:AB_141698 |
| Goat Anti-Rabbit IgG (H+L) Antibody, Alexa Fluor 594 Conjugated | Molecular Probes | Cat# A-11012, RRID:AB_141359 |
| Goat Anti-Mouse IgG1 Antibody, Alexa Fluor 488 Conjugated | Molecular Probes | Cat# A-21121, RRID:AB_141514 |
| Peroxidase-AffiniPure Goat Anti-Mouse IgG (H+L) antibody | Jackson ImmunoResearch Labs | Cat# 115-035-146, RRID:AB_2307392 |
| Peroxidase-AffiniPure Goat Anti-Rabbit IgG (H+L) antibody | Jackson ImmunoResearch Labs | Cat# 111-035-003, RRID:AB_2313567 |
| Donkey Anti-Rabbit IgG (H+L) Antibody, Alexa Fluor 488 Conjugated | Molecular Probes | Cat# A-21206, RRID:AB_141708 |
| Donkey Anti-Mouse IgG (H+L) Polyclonal Antibody, Alexa Fluor 647 Conjugated | Molecular Probes | Cat# A-31571, RRID:AB_162542 |
| anti Biotin | Jackson ImmunoResearch Labs | Cat# 200-002-211, RRID: AB_2339006 |

### Chemicals, Peptides, and Recombinant Proteins

| REAGENT | SOURCE | IDENTIFIER |
| --- | --- | --- |
| Proteinase K | Qiagen | Cat# 19133 |
| LumiGLO | Cell Signaling Technology | Cat# 7003S |
| OPTI-MEM media | Gibco | Cat# 31985070 |
| Lipofectamin 2000 | Invitrogen | Cat# 11668019 |
| Lipofectamin RNAiMAX | Invitrogen | Cat# 13778150 |
| DAPI | Sigma | Cat# 32670 |
| Mounting media | DAKO | Cat# S3023 |
| Streptavidin-sepharose beads | GE | Cat# GE17-5113-01 |
| Trypsin | Sigma-Aldrich | Cat# T6567 |
| Tetracycline hydrochloride | Sigma | Cat# T7660 |
| Biotin | Sigma | Cat# B4639 |
| Hygromycin B | Wisent | Cat# 450-141-XL |
| Puromycin | Wisent | Cat# 450-162-XL |
| Chloroquine | Sigma | Cat# C6628 |
| Polybrene | Aldrich | Cat# 10768 |
| Protease Inhibitor Cocktail | Sigma | Cat# P8340 |
| Critical Commercial Assays |  |  |
| Gateway™ BP Clonase™ II Enzyme mix | Invitrogen | Cat# 11789020 |
| Gateway™ LR Clonase™ II Enzyme mix | Invitrogen | Cat# 11791020 |
| QuikChange Lightning Site-Directed Mutagenesis Kit | Stratagene | Cat# 210518 |

### Deposited Data

| RESOURCE | SOURCE | IDENTIFIER |
| --- | --- | --- |
| Analyzed data | this study | http://prohits-web.lunenfeld.ca |
| Mitocarta 2.0 | PMID: 26450961 | PMID: 26450961 |
| Integrated Mitochondrial Protein Index (IMPI) | PMID: 26432830 | http://www.mrc-mbu.cam.ac.uk/impi |
| Gene ontology, GO enrichment analysis | PMID:12952881 | http://www.pantherdb.org |
| IMS APEX proteomics data | PMID: 25002142 | PMID: 25002142 |
| Matrix APEX proteomics data | PMID: 23371551 | PMID: 23371551 |
| Raw and analyzed data | this study | MassIVE # |
| Subcellular location data version 18.1 | The Human Protein Atlas | HPA |

### Experimental Models: Cell Lines

| REAGENT | SOURCE | IDENTIFIER |
| --- | --- | --- |
| control human fibroblasts | Montreal Children Hospital cellbank |  |
| Flp-In™ T-REx™ 293 | Invitrogen | Cat# R78007 |
| Phoenix | a gift from Garry P. Nolan |  |

### Oligonucleotides

| REAGENT | SOURCE | IDENTIFIER |
| --- | --- | --- |
| Primers and clones used | this study | Table S8 |
| EXD2 siRNA | Dharmacon | Cat# L-020899-02-0005 |

### Recombinant DNA

| REAGENT | SOURCE | IDENTIFIER |
| --- | --- | --- |
| pDEST-pcDNA5-BirA*-FLAG C-term | in house | PMID: 24255178 |
| pDEST-pcDNA5-BirA*-FLAG-GFP | in house | PMID: 24255178 |
| pOG44 | Invitrogen | Cat# V600520 |
| mCherry-ER-3 | Addgene | Cat# 55041 |
| Gateway™ pDONR™221 | Invitrogen | Cat# 12536017 |
| pDEST-pcDNA5-BirA*-FLAG N-term | in house | PMID: 24255178 |
| pBabe-gtw-rfA | in house | PMID: 16632485 |

### Software and Algorithms

| RESOURCE | SOURCE | IDENTIFIER |
| --- | --- | --- |
| ProHits-viz analysis | PMID: 28661499 | https://prohits-viz.lunenfeld.ca<br>PMID: 28661499 |
| Cytoscape | PMID: 14597658 | https://cytoscape.org<br>PMID: 14597658 |
| Fiji | PMID: 22743772 | PMID: 22743772 |
| PDB Viewer | PMID: 9504803 | http://www.expasy.org/spdbv/<br>PMID: 9504803 |
| TMHMM transmembrane prediction software | PMID: 11152613 | www.cbs.dtu.dk/services/TMHMM/<br>PMID: 11152613 |
| Prism 8 | GraphPad | https://www.graphpad.com |
| Venny 2.1 | Venny | https://bioinfogp.cnb.csic.es/tools/venny/index.html |
| BioVenn | BioVenn | http://www.biovenn.nl<br>PMID: 18925949 |

