## Supplementary Figures for "A high-density human mitochondrial proximity interaction network"

### Figure S1

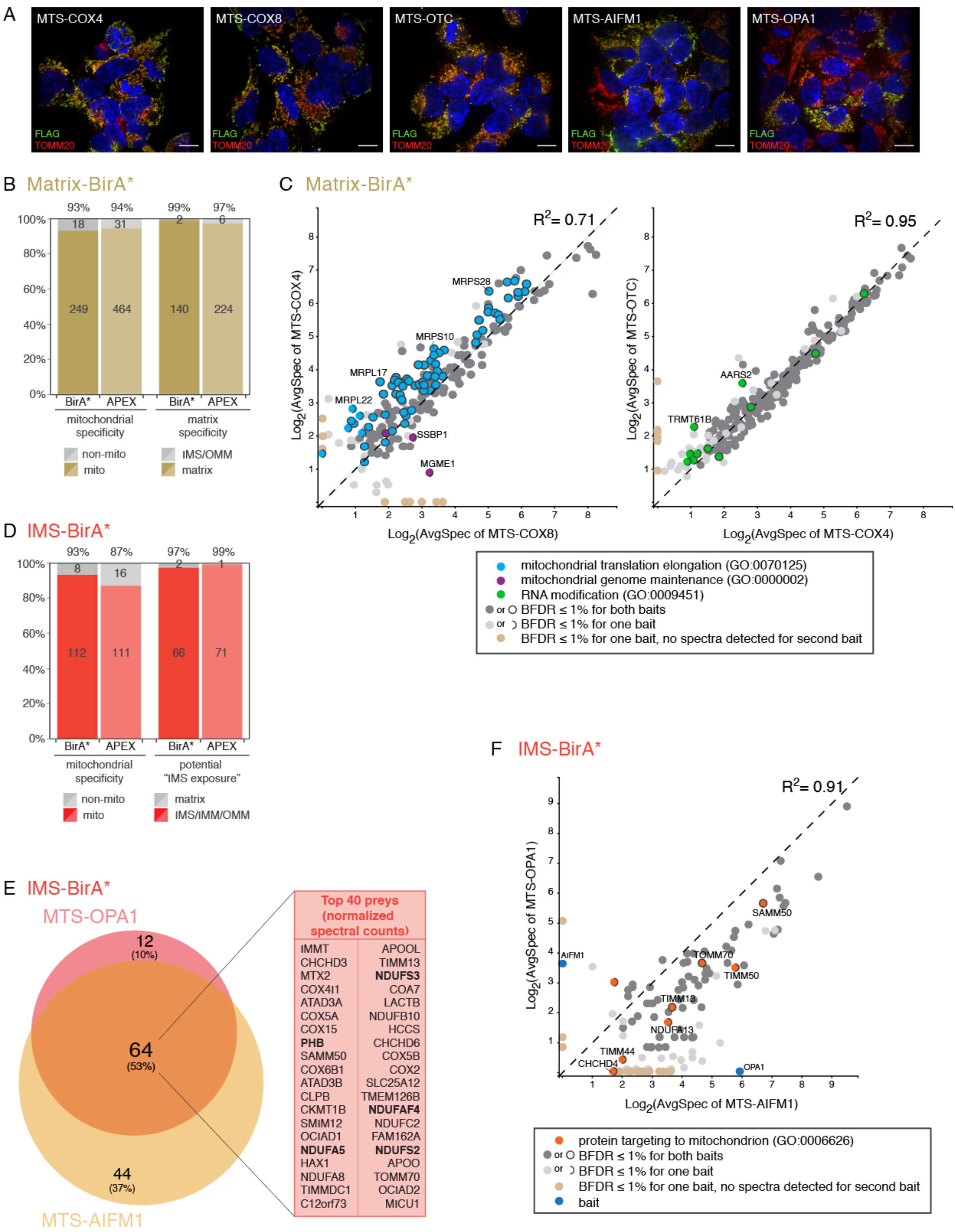
A
MTS-COX4
MTS-COX8
MTS-OTC
MTS-AIFM1
MTS-OPA1
FLAG
TOMM20
B Matrix-BirA*
93% 94% 99% 97%
18 31 2 6
249 464 140 224
100% 80% 60% 40% 20% 0%
BirA* APEX BirA* APEX
mitochondrial specificity
matrix specificity
non-mito
mito
IMS/OMM
matrix
C Matrix-BirA*
$R^2$= 0.71
$R^2$= 0.95
MRPS28
MRPS10
MRPL17
MRPL22
SSBP1
MGME1
AARS2
TRMT61B
Log2(AvgSpec of MTS-COX4)
Log2(AvgSpec of MTS-COX8)
Log2(AvgSpec of MTS-OTC)
Log2(AvgSpec of MTS-COX4)
mitochondrial translation elongation (GO:0070125)
mitochondrial genome maintenance (GO:0000002)
RNA modification (GO:0009451)
or BFDR ≤ 1% for both baits
or BFDR ≤ 1% for one bait
BFDR ≤ 1% for one bait, no spectra detected for second bait
D IMS-BirA*
93% 87% 97% 99%
8 16 2 1
112 111 66 71
100% 80% 60% 40% 20% 0%
BirA* APEX BirA* APEX
mitochondrial specificity
potential "IMS exposure"
non-mito
mito
matrix
IMS/IMM/OMM
E IMS-BirA*
MTS-OPA1
12 (10%)
64 (53%)
44 (37%)
MTS-AIFM1
Top 40 preys (normalized spectral counts)
IMMT APOOL
CHCHD3 TIMM13
MTX2 NDUFS3
COX4I1 COA7
ATAD3A LACTB
COX5A NDUFB10
COX15 HCCS
PHB CHCHD6
SAMM50 COX5B
COX6B1 COX2
ATAD3B SLC25A12
CLPB TMEM126B
CKMT1B NDUFAF4
SMIM12 NDUFC2
OCIAD1 FAM162A
NDUFA5 NDUFS2
HAX1 APOO
NDUFA8 TOMM70
TIMMDC1 OCIAD2
C12orf73 MICU1
F IMS-BirA*
$R^2$= 0.91
SAMM50
AIFM1
TOMM70
TIMM50
TIMM13
NDUFA13
TIMM44
CHCHD4
OPA1
Log2(AvgSpec of MTS-OPA1)
Log2(AvgSpec of MTS-AIFM1)
protein targeting to mitochondrion (GO:0006626)
or BFDR ≤ 1% for both baits
or BFDR ≤ 1% for one bait
BFDR ≤ 1% for one bait, no spectra detected for second bait
bait

### Figure S2

A

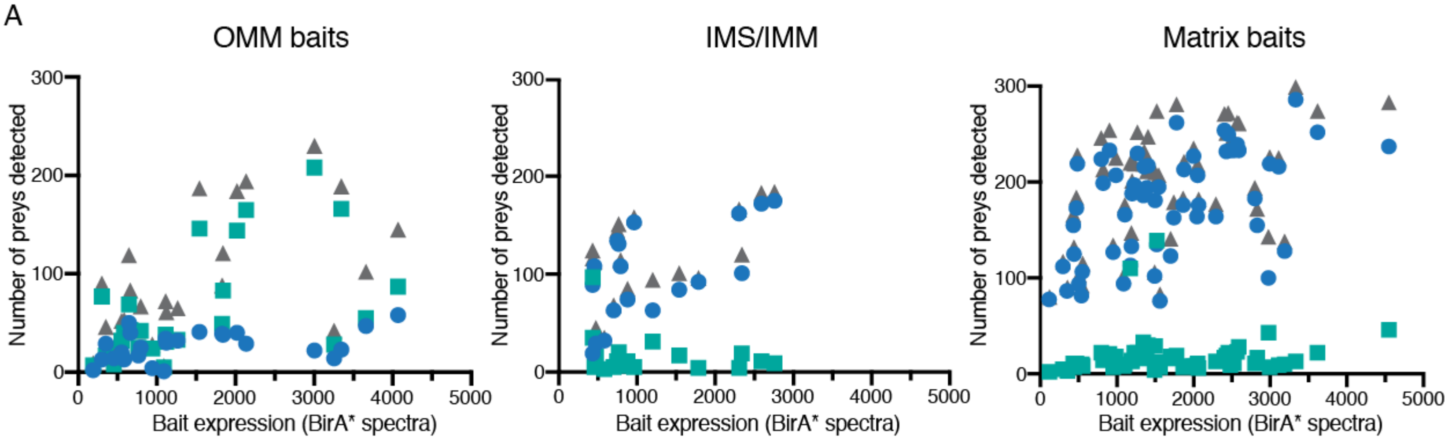


| | OMM | IMS/IMM | Matrix |
|---|---|---|---|
| ● Mitocarta preys | 0.4269, P=0.0333 | 0.6313, P=0.0050 | 0.4929, P<0.0001 |
| ■ Non-Mitocarta preys | 0.5436, P=0.0050 | -0.2699, n.s. | 0.1060, n.s |
| ▲ Total | 0.5876, P=0.0020 | 0.5718, P=0.0132 | 0.5007, P<0.0001 |

B

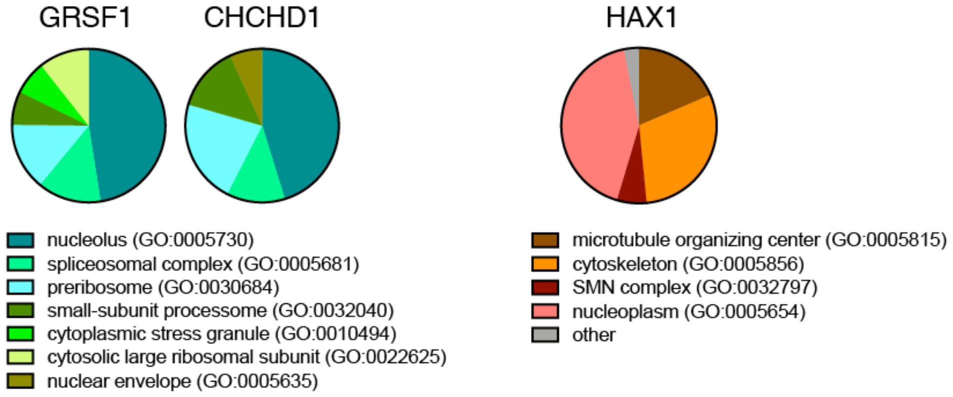


C

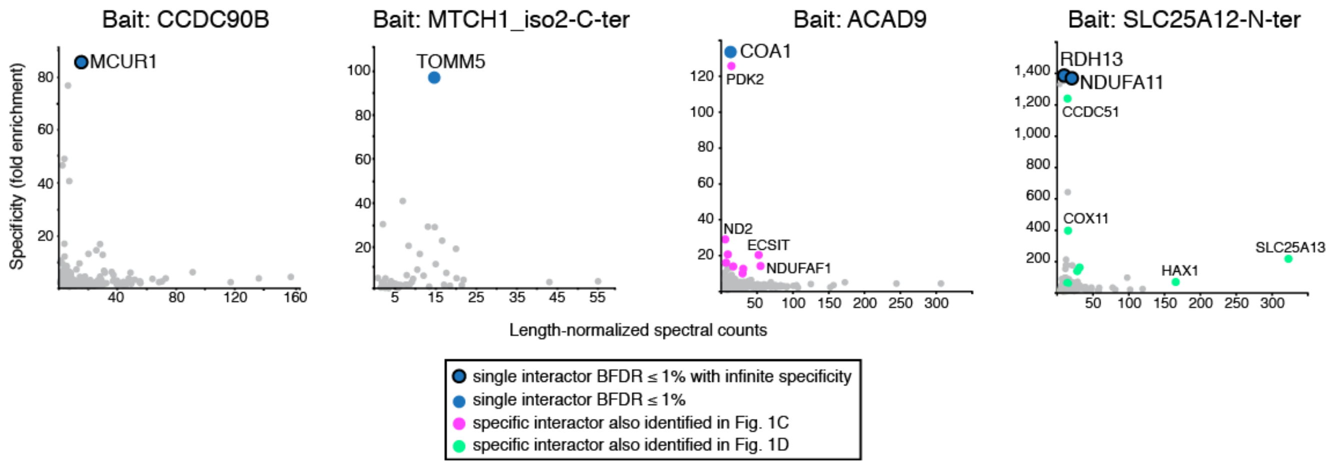

### Figure S3

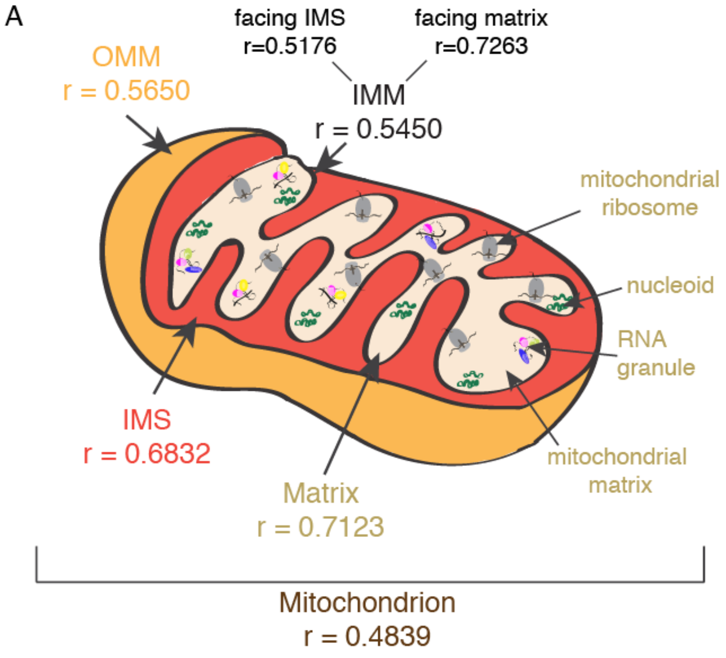


OMM ↔ matrix
r = 0.2812

OMM ↔ IMM
r = 0.3394

matrix ↔ IMM
r = 0.4201

| | IMM facing IMS | IMM facing matrix |
|---|---|---|
| OMM | 0.4173 | 0.3025 |
| matrix | 0.3131 | 0.5365 |

| Matrix GOCC term | RNP granule | mitochondrial ribosome | mitochondrial matrix |
|---|---|---|---|
| RNP granule | 0.6532 | 0.6567 | 0.6403 |
| mitochondrial ribosome | 0.6567 | 0.7715 | 0.7604 |
| mitochondrial matrix | 0.6403 | 0.7604 | 0.7962 |

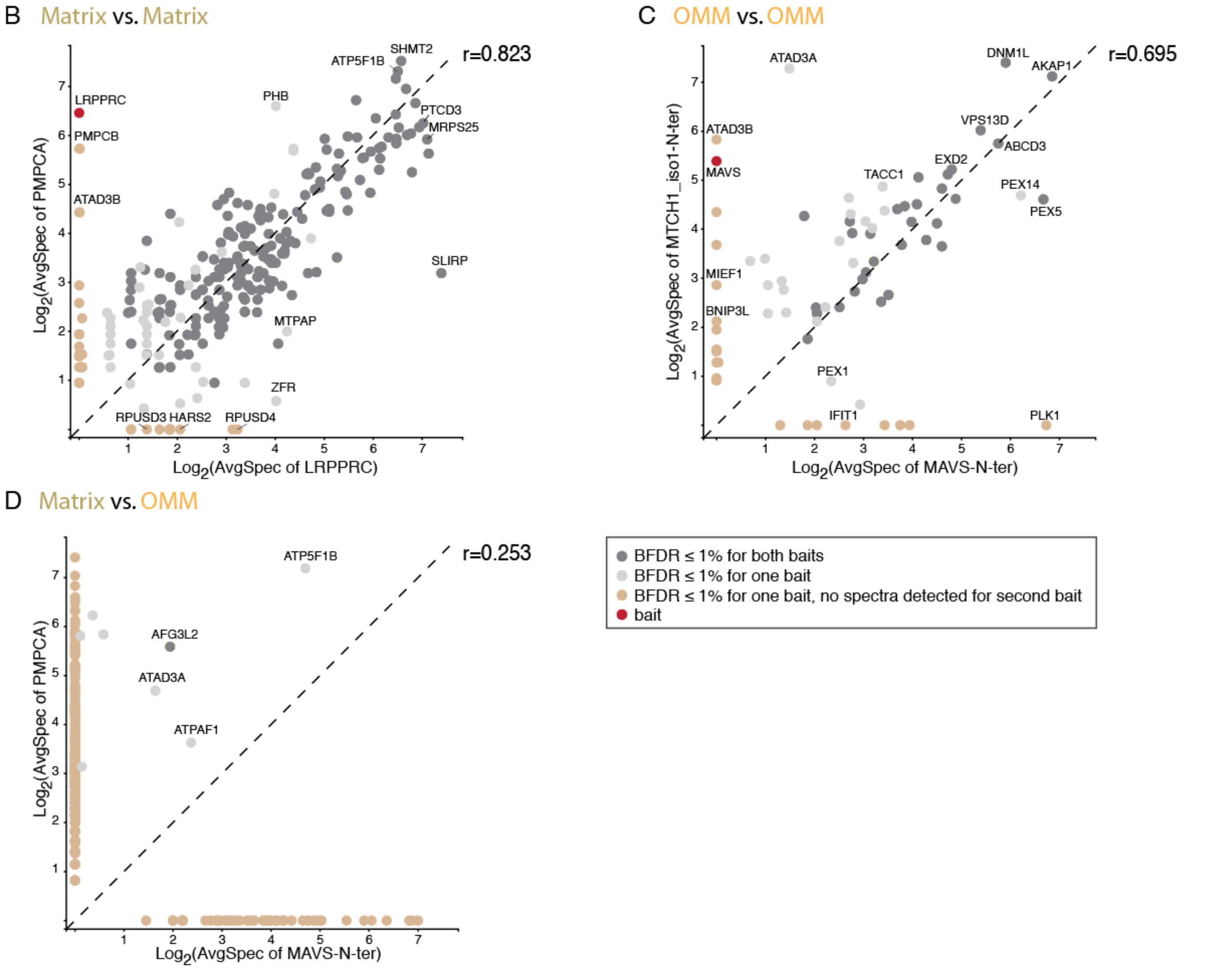

### Figure S4

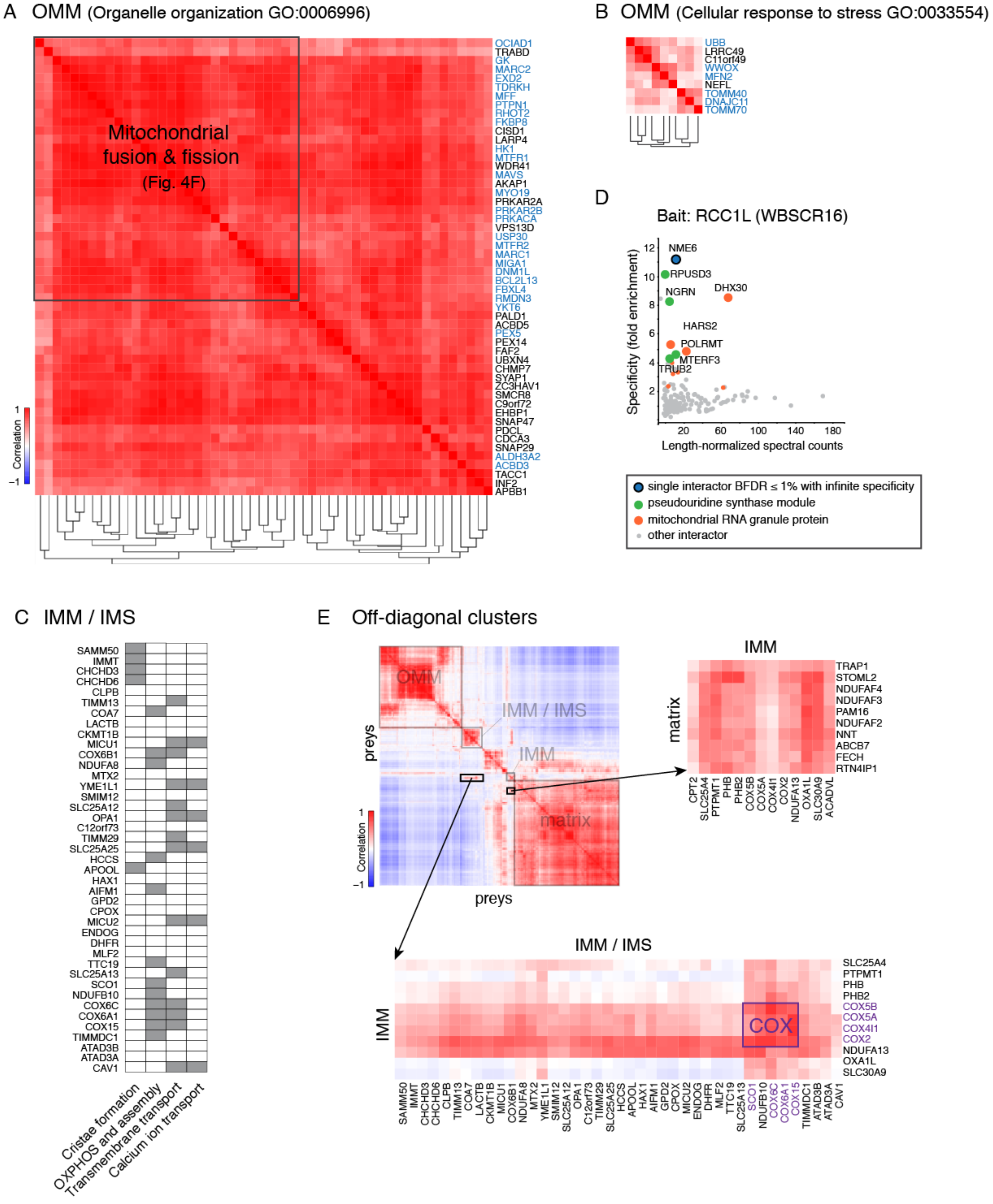
A OMM (Organelle organization GO:0006996)
Mitochondrial fusion & fission (Fig. 4F)
OCIAD1
TRABD
GK
MARC2
EXD2
TDRKH
MFF
PTPN1
RHOT2
FKBP8
CISD1
LARP4
HK1
MTFR1
WDR41
MAVS
AKAP1
MYO19
PRKAR2A
PRKAR2B
PRKACA
VPS13D
USP30
MTFR2
MARC1
MIGA1
DNM1L
BCL2L13
FBXL4
RMDN3
YKT6
PALD1
ACBD5
PEX5
PEX14
FAF2
UBXN4
CHMP7
SYAP1
ZC3HAV1
SMCR8
C9orf72
EHBP1
SNAP47
PDCL
CDCA3
SNAP29
ALDH3A2
ACBD3
TACC1
INF2
APBB1
Correlation
1
−1
B OMM (Cellular response to stress GO:0033554)
UBB
LRRC49
C11orf49
WWOX
MFN2
NEFL
TOMM40
DNAJC11
TOMM70
D
Bait: RCC1L (WBSCR16)
Specificity (fold enrichment)
Length-normalized spectral counts
NME6
RPUSD3
NGRN
DHX30
HARS2
POLRMT
MTERF3
TRUB2
single interactor BFDR ≤ 1% with infinite specificity
pseudouridine synthase module
mitochondrial RNA granule protein
other interactor
C IMM / IMS
SAMM50
IMMT
CHCHD3
CHCHD6
CLPB
TIMM13
COA7
LACTB
CKMT1B
MICU1
COX6B1
NDUFA8
MTX2
YME1L1
SMIM12
SLC25A12
OPA1
C12orf73
TIMM29
SLC25A25
HCCS
APOOL
HAX1
AIFM1
GPD2
CPOX
MICU2
ENDOG
DHFR
MLF2
TTC19
SLC25A13
SCO1
NDUFB10
COX6C
COX6A1
COX15
TIMMDC1
ATAD3B
ATAD3A
CAV1
Cristae formation
OXPHOS and assembly
Transmembrane transport
Calcium ion transport
E Off-diagonal clusters
OMM
IMM / IMS
IMM
matrix
preys
Correlation
IMM
matrix
TRAP1
STOML2
NDUFAF4
NDUFAF3
PAM16
NDUFAF2
NNT
ABCB7
FECH
RTN4IP1
CPT2
SLC25A4
PTPMT1
PHB
PHB2
COX5B
COX5A
COX4I1
COX2
NDUFA13
OXA1L
SLC30A9
ACADVL
IMM / IMS
IMM
COX
SLC25A4
PTPMT1
PHB
PHB2
COX5B
COX5A
COX4I1
COX2
NDUFA13
OXA1L
SLC30A9

### Figure S5

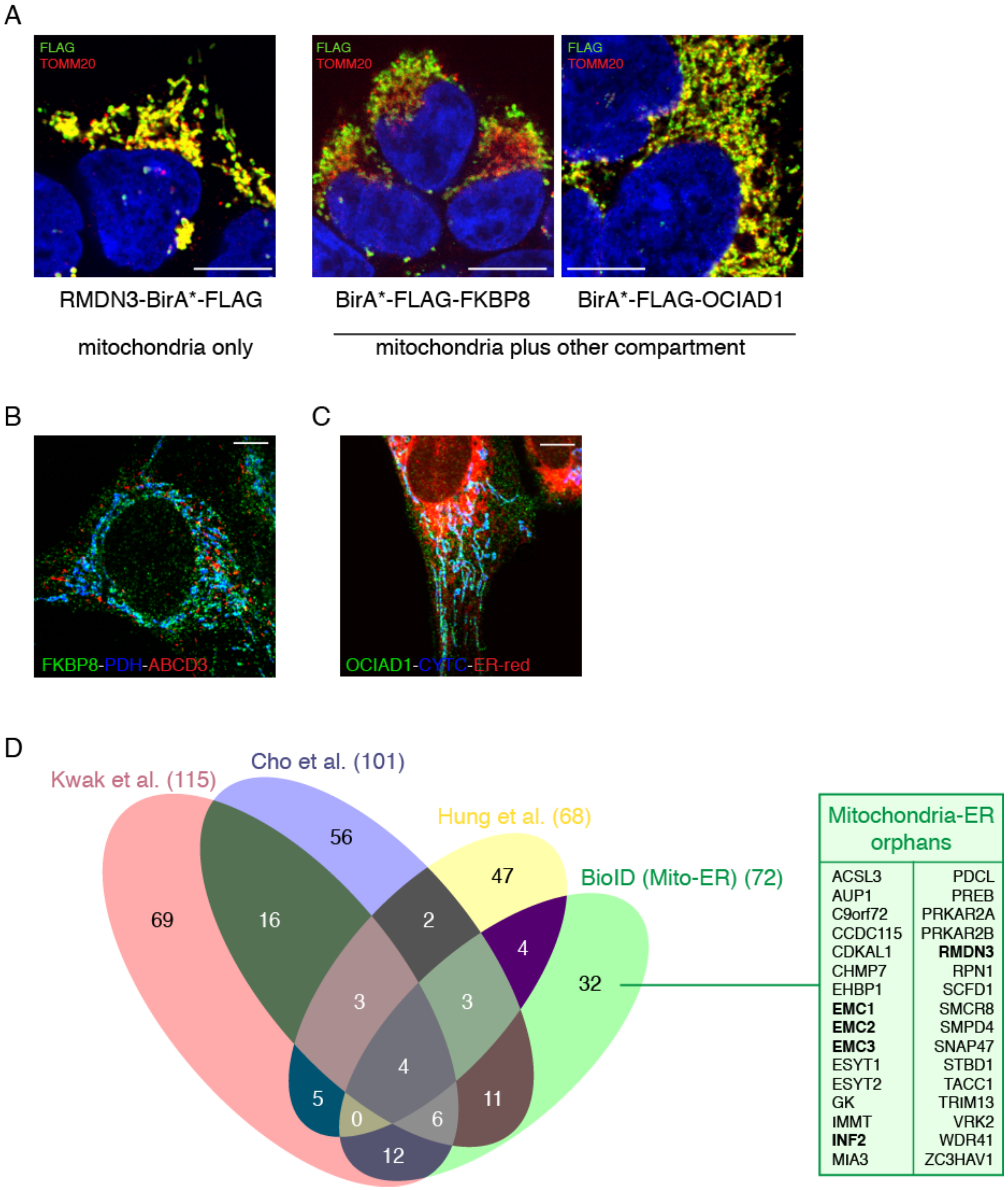
A
FLAG
TOMM20
RMDN3-BirA*-FLAG
BirA*-FLAG-FKBP8
BirA*-FLAG-OCIAD1
mitochondria only
mitochondria plus other compartment
B
FKBP8-PDH-ABCD3
C
OCIAD1-CYTC-ER-red
D
Kwak et al. (115)
Cho et al. (101)
Hung et al. (68)
BioID (Mito-ER) (72)
69
16
56
2
47
4
32
3
3
4
5
0
6
11
12
Mitochondria-ER orphans
ACSL3
AUP1
C9orf72
CCDC115
CDKAL1
CHMP7
EHBP1
EMC1
EMC2
EMC3
ESYT1
ESYT2
GK
IMMT
INF2
MIA3
PDCL
PREB
PRKAR2A
PRKAR2B
RMDN3
RPN1
SCFD1
SMCR8
SMPD4
SNAP47
STBD1
TACC1
TRIM13
VRK2
WDR41
ZC3HAV1

### Figure S6

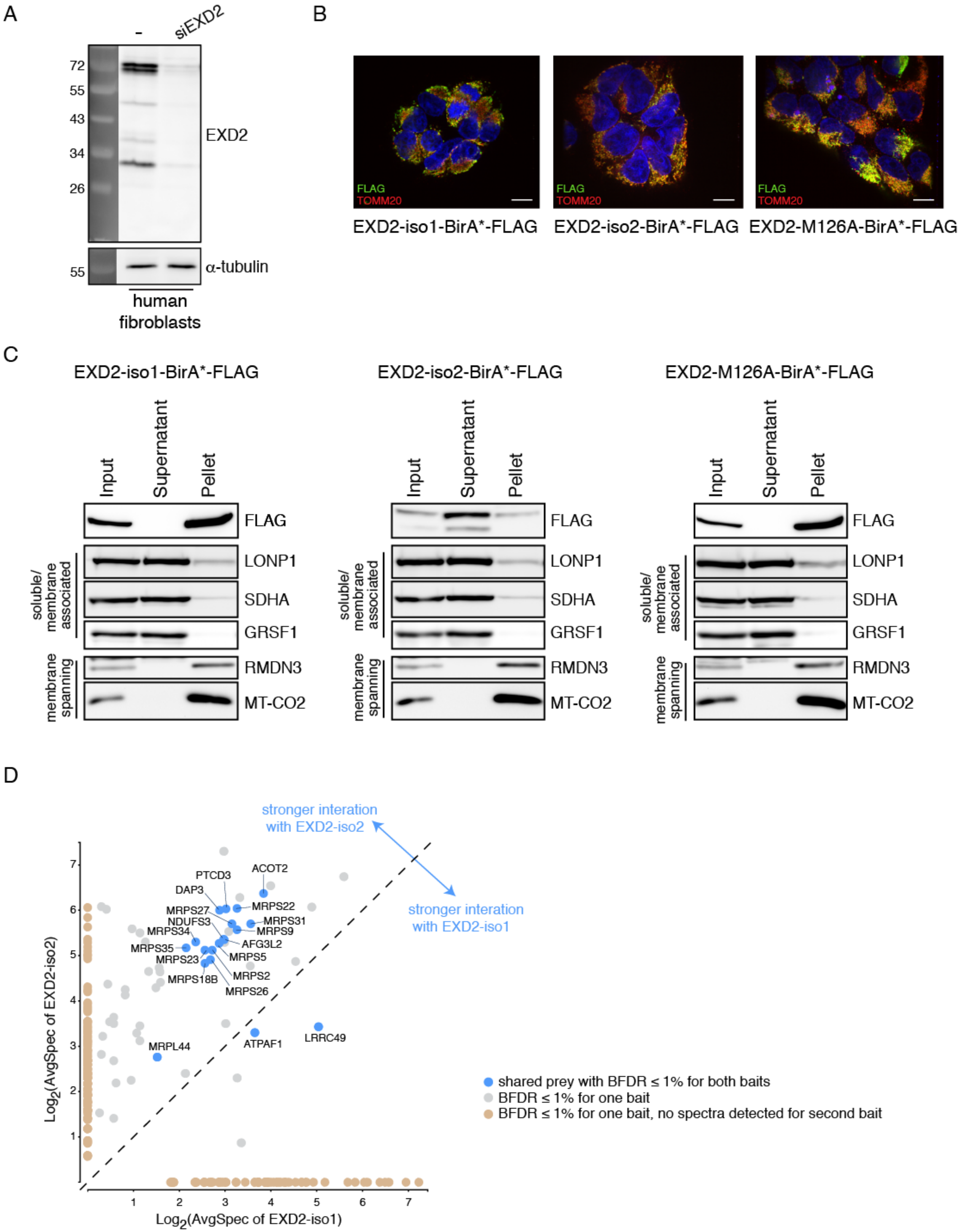
A
-
siEXD2
72
55
43
34
26
EXD2
55
α-tubulin
human fibroblasts
B
FLAG
TOMM20
EXD2-iso1-BirA*-FLAG
EXD2-iso2-BirA*-FLAG
EXD2-M126A-BirA*-FLAG
C
EXD2-iso1-BirA*-FLAG
EXD2-iso2-BirA*-FLAG
EXD2-M126A-BirA*-FLAG
Input
Supernatant
Pellet
FLAG
LONP1
SDHA
GRSF1
RMDN3
MT-CO2
soluble/ membrane associated
membrane spanning
D
stronger interation with EXD2-iso2
stronger interation with EXD2-iso1
ACOT2
PTCD3
DAP3
MRPS27
MRPS22
NDUFS3
MRPS31
MRPS34
MRPS9
AFG3L2
MRPS35
MRPS5
MRPS23
MRPS2
MRPS18B
MRPS26
MRPL44
ATPAF1
LRRC49
Log2(AvgSpec of EXD2-iso2)
Log2(AvgSpec of EXD2-iso1)
shared prey with BFDR ≤ 1% for both baits
BFDR ≤ 1% for one bait
BFDR ≤ 1% for one bait, no spectra detected for second bait

### Figure S7

A

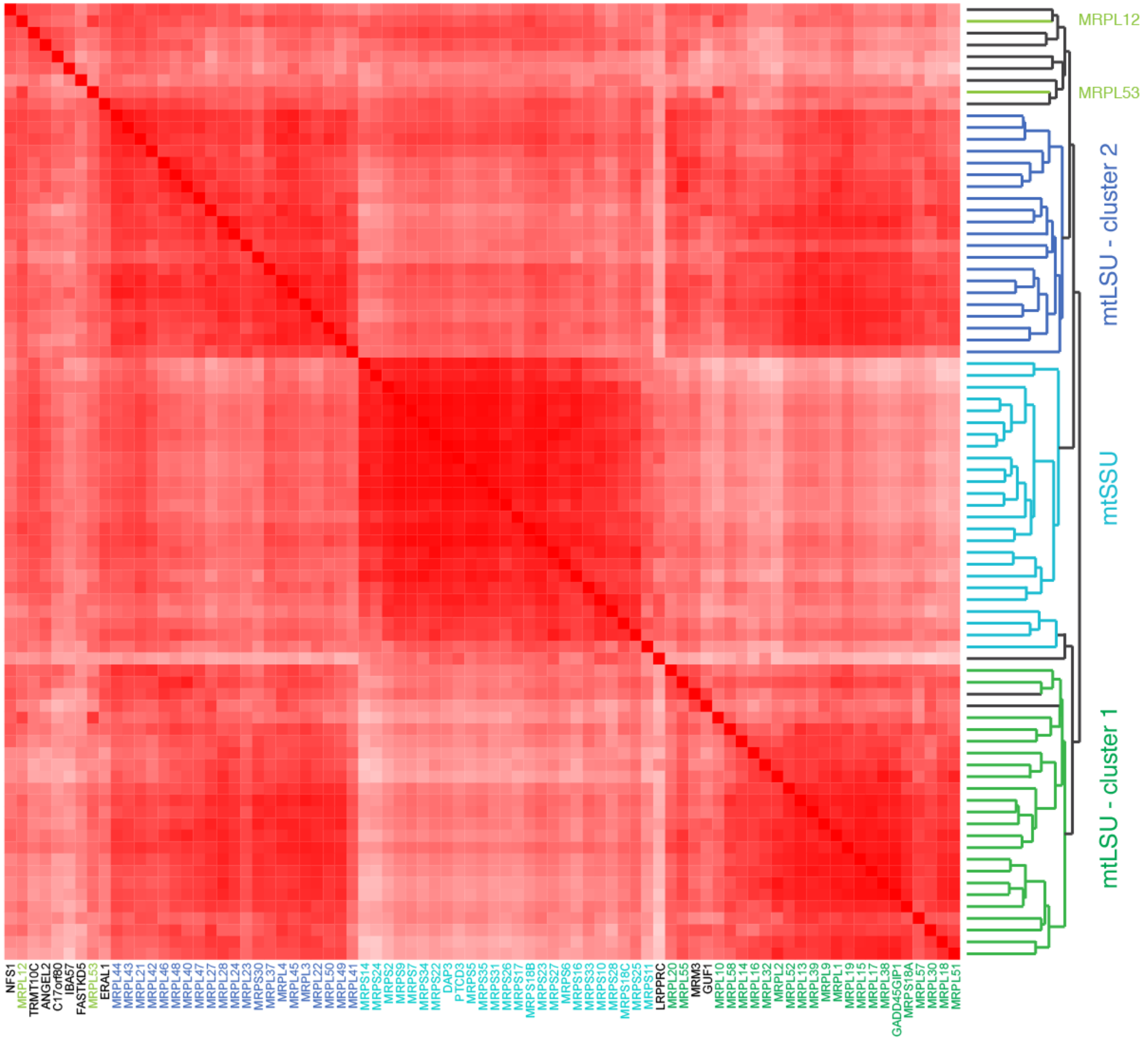


B mtLSU - cluster 3

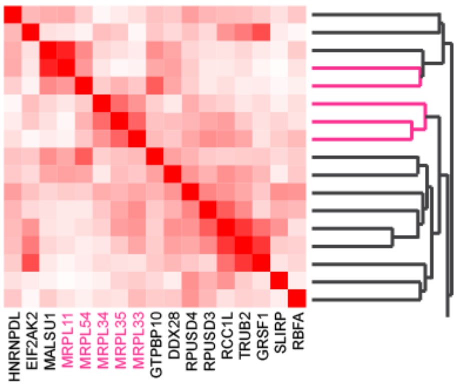
